# Trustworthy Sleep Staging from EEG: Deep Ensembles, MC Dropout, and Predictive Calibration

**DOI:** 10.1101/2025.08.23.671918

**Authors:** BC Samiksha, Dana Vrajitoru

## Abstract

Reliable sleep stage classification from EEG signals is critical for the development of clinical decision support systems. However, many deep learning models lack mechanisms for estimating predictive uncertainty, which is important for trust and interpretability. This work explores the use of Monte Carlo Dropout and Deep Ensembles to estimate uncertainty in automatic sleep staging. We apply these methods to convolutional and recurrent neural architectures trained on a publicly available EEG dataset. Evaluation includes calibration analysis and visualization of predictive entropy across sleep stages. LSTM-based models demonstrated more consistent trustworthiness and calibration across runs, while EEGNet remained appealing for lightweight deployment.The results highlight the value of incorporating uncertainty estimates to improve model transparency and support safe use in medical contexts.

## 1 Introduction

Accurate sleep stage classification from EEG signals is essential for diagnosing sleep disorders, conducting cognitive studies, and supporting long-term health monitoring. Manual scoring is labor-intensive and subject to inconsistencies, which has led to the growing adoption of deep learning models for automating this task. While many of these models achieve strong performance, they often lack the ability to express how confident they are in their predictions. This omission is particularly problematic in clinical settings, where model outputs are used to support or guide human decision-making. In this work, we investigate the role of predictive uncertainty in EEG-based sleep staging by integrating Monte Carlo Dropout and Deep Ensembles into two representative architectures: EEGNet and an LSTM with temporal attention. Our goal is to move beyond accuracy alone, and toward calibrated, interpretable, and trustworthy model behavior.

## 2 Related Work

Deep learning has become the standard approach for automated sleep staging from EEG. Convolutional models such as EEGNet have been widely adopted due to their efficiency and strong performance on dual-channel data [1]. Recurrent neural networks, including LSTMs, have also shown promise in capturing temporal dependencies across sleep epochs [2]. Hybrid models incorporating attention mechanisms have recently been explored to improve interpretability and context modeling [3]. However, these approaches typically optimize for accuracy alone and do not quantify predictive uncertainty.

## 3 Model Architectures

We propose two temporal deep learning architectures for EEG-based sleep staging: EEGNet with temporal attention and an LSTM-attention hybrid. EEGNet leverages Conv1D, DepthwiseConv1D, and SeparableConv1D layers to efficiently extract spatial and temporal EEG features with minimal parameters. The LSTM model encodes long-range temporal dependencies and enhances feature representation via multi-head temporal self-attention. Both architectures incorporate dropout regularization and max-norm constraints, and output stage probabilities through a softmax classifier. For uncertainty estimation, we apply Monte Carlo Dropout and Deep Ensembles during inference to quantify predictive confidence.

### 3.1 EEGNet Architecture

The architecture begins with a temporal Conv1D layer followed by DepthwiseConv1D and SeparableConv1D, reflecting spatial and frequency filtering. Each convolution is followed by batch normalization, ELU activation, max-pooling, and dropout. A multi-head attention block is inserted before global average pooling and a dense softmax classifier.

#### Algorithm 1

EEGNet with Temporal Attention

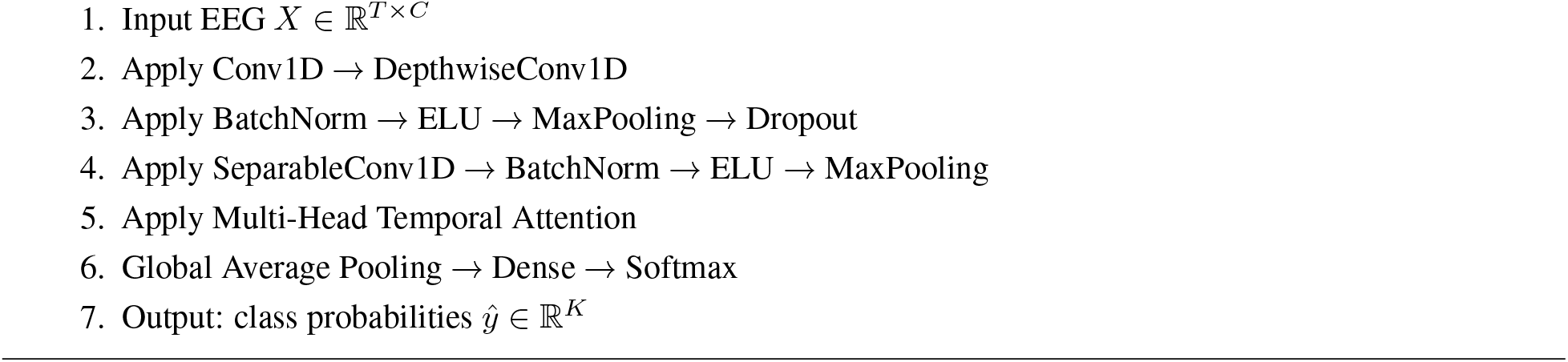

### 3.2 LSTM Architecture

This model uses a single bidirectional LSTM layer with 128 units, followed by DepthwiseConv1D, SeparableConv1D, and a multi-head temporal attention mechanism. Like EEGNet, it concludes with average pooling and a dense softmax layer. Both models apply max-norm kernel constraints to control weight magnitudes and use ReLU or ELU activations with dropout for regularization.

#### Algorithm 2

LSTM with Temporal Attention

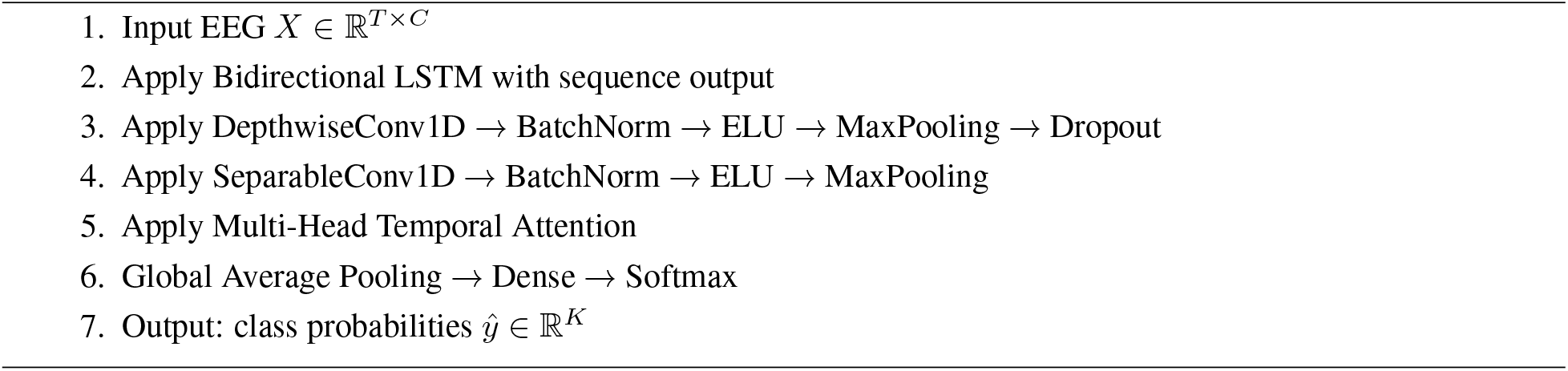

### 3.3 Uncertainty Estimation Approaches

We investigate two uncertainty estimation methods:

#### Monte Carlo Dropout

In this approach, dropout layers remain active during inference to approximate Bayesian posterior sampling. We perform *M* = 20 stochastic forward passes and compute the predictive mean as:

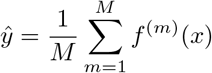

Uncertainty is quantified using the predictive entropy:

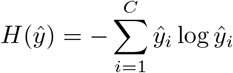

where *C* is the number of sleep stage classes and *ŷ*_*i*_ is the averaged probability for class *i*. This captures the model’s confidence in its predictions across multiple forward passes.

#### Deep Ensembles

We construct an ensemble of *K* = 5 independently trained models, each initialized with a unique random seed and trained on the same data distribution. For an input *x*, each model *f*_*k*_(*x*) outputs a softmax probability vector *p*_*k*_ ∈ ℝ^*C*^, where *C* = 5 is the number of sleep stages. The final predictive distribution is obtained by averaging across the ensemble:

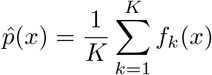

Uncertainty is quantified via the empirical standard deviation of the ensemble predictions:

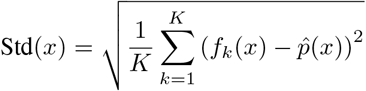

### 3.4 Training and Evaluation Setup

All models are trained using the Adam optimizer with learning rate *α* = 5 *×* 10^−4^ and batch size *B* = 16. The categorical cross-entropy loss function is minimized over the training set 𝒟_train_:

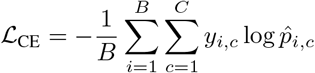

where *y*_*i,c*_ is the one-hot ground truth and 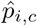 is the predicted probability for class *c* on sample *i*.

To mitigate overfitting, we apply early stopping with a patience of 3 epochs based on validation loss. Training is capped at 20 epochs. To address class imbalance, we compute per-class sample weights:

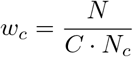

where *N* is the total number of training examples and *N*_*c*_ is the number of samples in class *c*. These weights are applied

to each loss term during backpropagation.

Data is split using stratified sampling on a per-subject-night basis to preserve class distribution. All experiments are conducted using mixed-precision training with the ‘mixed float16’ policy in TensorFlow to accelerate convergence.

### 3.5 Evaluation Metrics

Performance is evaluated using the following metrics:

- **Accuracy**: Standard classification accuracy across all test samples.
- **Negative Log-Likelihood (NLL)**:

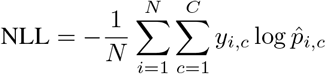
- **Brier Score**:

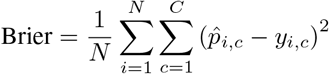
- **Expected Calibration Error (ECE):** Computed using *B* = 15 confidence bins as:

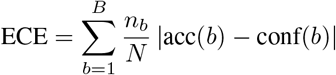

where *n*_*b*_ is the number of samples in bin *b*, acc(*b*) is accuracy in the bin, and conf(*b*) is the average confidence.

All metrics are computed on the held-out test set. Accuracy captures overall classification performance, while NLL penalizes overconfident incorrect predictions. The Brier score measures the squared error between predicted probabilities and true labels. ECE estimates how well predicted confidence aligns with actual accuracy using 15 confidence bins. Together, these metrics provide a comprehensive view of both prediction quality and calibration.

## 4 Experiments and Results

### 4.1 Performance Metrics

We evaluate four model configurations: EEGNet and LSTM, each paired with either Monte Carlo Dropout (MC Dropout) or Deep Ensembles. Models are trained on the Sleep-EDF Expanded dataset and evaluated using subject-night-level stratified test splits. To assess generalization, we report overall classification accuracy and supplement it with class-wise precision, recall, and F1-score.

**Table 1.** Macro-averaged test performance metrics across all evaluated models.

| Model | Acc | Prec | Rec | F1 |
| --- | --- | --- | --- | --- |
| EEGNet MC | 0.8943 | 0.7390 | 0.8285 | 0.7669 |
| EEGNet Ensemble | 0.8716 | 0.7020 | 0.8123 | 0.7367 |
| LSTM MC | 0.8952 | 0.7161 | 0.7827 | 0.7582 |
| LSTM Ensemble | 0.8891 | 0.7381 | 0.7824 | 0.7606 |

Let 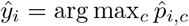 be the predicted class for sample *i*, and *y*_*i*_ the true class. Overall accuracy is computed as:

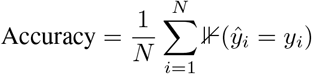

Per-class performance is measured as:

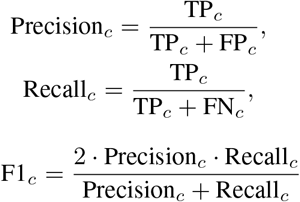

where TP_*c*_, FP_*c*_, and FN_*c*_ are true positives, false positives, and false negatives for class *c*.

LSTM with MC Dropout achieves the highest overall accuracy, while EEGNet with Deep Ensembles demonstrates the most stable class-wise performance, particularly for difficult stages such as N1 and REM.

## 5 Conclusion

We presented a comparative evaluation of uncertainty-aware deep learning models for automatic sleep stage classification using dual-channel EEG (Fpz-Cz and Pz-Oz) from the Sleep-EDF dataset. Our study assessed both classification performance and predictive reliability across two representative architectures: EEGNet and LSTM, each extended with Monte Carlo Dropout and Deep Ensembles to quantify uncertainty.

LSTM models generally yielded more consistent accuracy and calibration, particularly when combined with MC Dropout, making them strong candidates for uncertainty-sensitive applications. EEGNet, while slightly less accurate, showed improved calibration under Deep Ensembles and remains favorable for resource-constrained deployment due to its compact design.

These findings underscore the need to balance performance with trustworthiness when designing clinical decision support tools. Incorporating uncertainty-aware training and evaluation can help ensure safer deployment and greater interpretability. Future work will explore multimodal PSG inputs, subject-specific fine-tuning, and selective abstention mechanisms that defer uncertain predictions to human experts.

## Supporting information

main.py

## Notes

https://github.com/sameekshya1999/Trustworthy-Sleep-Staging/tree/master

### Competing Interest Statement

The authors have declared no competing interest.

