## Supplementary material for "Trustworthy Sleep Staging from EEG: Deep Ensembles, MC Dropout, and Predictive Calibration": main.py: main.py

```
import numpy as np
import tensorflow as tf
from tensorflow.keras import layers, models, constraints
from tensorflow.keras.utils import Sequence
from tensorflow.keras.callbacks import EarlyStopping
import mne
import urllib.request
import os
from sklearn.model_selection import train_test_split
from sklearn.utils.class_weight import compute_class_weight
from sklearn.metrics import precision_recall_fscore_support, confusion_matrix
from concurrent.futures import ThreadPoolExecutor
from tqdm import tqdm
import warnings
import matplotlib.pyplot as plt
import seaborn as sns
import sklearn
import gc
import random
import pandas as pd

warnings.filterwarnings("ignore", message="Channels contain different highpass filters")
warnings.filterwarnings("ignore", message="Channels contain different lowpass filters")
warnings.filterwarnings("ignore", message="Highpass cutoff frequency")

np.random.seed(42)
tf.random.set_seed(42)
random.seed(42)
os.environ['PYTHONHASHSEED'] = '42'

from tensorflow.keras.mixed_precision import set_global_policy
set_global_policy('mixed_float16')

warnings.filterwarnings("ignore", category=DeprecationWarning)
mne.set_log_level('ERROR')

NUM_SUBJECTS = 20
NUM_NIGHTS = 2
BASE_URL = "https://physionet.org/files/sleep-edfx/1.0.0/"
TARGET_CHANNELS = ['EEG Fpz-Cz', 'EEG Pz-Oz']
EPOCH_DURATION = 30
BATCH_SIZE = 16
EPOCHS = 20
SAMPLING_RATE = 50
TELEMETRY_SUBJECTS = [2, 4, 5, 6, 7, 12, 13]
POSSIBLE_HYPNO_LETTERS = 'CHJPUVAEMORW'
print(f"scikit-learn version: {sklearn.__version__}")

nb_classes = 5

def fetch_data(subject_id, night, record_type='PSG'):
    try:
        dataset_id = subject_id
        folder = "sleep-cassette" if night == 1 else "sleep-telemetry"
        if night == 1:
            prefix = f"SC4{dataset_id:02d}"
        else:
            if subject_id not in TELEMETRY_SUBJECTS:
                return None
            telemetry_map = {2: 702, 4: 704, 5: 705, 6: 706, 7: 707, 12: 712, 13: 713}
            prefix = f"ST{telemetry_map.get(subject_id, 700 + dataset_id)}"
        os.makedirs("sleep_edf", exist_ok=True)
        if record_type == 'PSG':
            base_suffix = "E" if night == 1 else "J"
            file_name = f"{prefix}{night if night == 1 else 2}{base_suffix}0-PSG.edf"
            url = f"{BASE_URL}{folder}/{file_name}"
            local_file = os.path.join("sleep_edf", file_name)
            if os.path.exists(local_file):
                return local_file
            urllib.request.urlretrieve(url, local_file)
            print(f"Downloaded {file_name}")
            return local_file
        else:
            base_suffix = "E" if night == 1 else "J"
            for letter in POSSIBLE_HYPNO_LETTERS:
                hypno_suffix = base_suffix + letter
                file_name = f"{prefix}{night if night == 1 else 2}{hypno_suffix}-Hypnogram.edf"
                url = f"{BASE_URL}{folder}/{file_name}"
                local_file = os.path.join("sleep_edf", file_name)
                if os.path.exists(local_file):
                    return local_file
                try:
                    urllib.request.urlretrieve(url, local_file)
                    print(f"Downloaded {file_name}")
                    return local_file
                except urllib.error.HTTPError as e:
                    if e.code != 404:
                        raise
            return None
    except urllib.error.HTTPError as e:
        print(f"HTTP Error {e.code} fetching {file_name if 'file_name' in locals() else 'file'}: {e.reason}")
        return None
    except Exception as e:
        print(f"Error fetching {file_name if 'file_name' in locals() else 'file'}: {e}")
        return None

def get_available_subjects():
    available = []
    with ThreadPoolExecutor(max_workers=10) as executor:
        futures = []
        for subject_id in range(NUM_SUBJECTS):
            for night in range(1, NUM_NIGHTS + 1):
                futures.append((
                    subject_id,
                    night,
                    executor.submit(
                        lambda s, n: (
                            fetch_data(s, n, 'PSG') is not None and
                            fetch_data(s, n, 'Hypnogram') is not None
                        ),
                        subject_id, night
                    )
                ))
        for subject_id, night, future in tqdm(futures, desc="Checking availability"):
            if future.result():
                available.append((subject_id, night))
    print(f"Available subject-night pairs: {available}")
    return available

def augment_data(X):
    noise = np.random.normal(0, 0.01, X.shape)
    shift = np.random.randint(-50, 50)
    X_aug = np.roll(X + noise, shift, axis=1)
    return X_aug

def process_subject_night(subject_id, night):
    try:
        psg_file = fetch_data(subject_id, night, 'PSG')
        hypno_file = fetch_data(subject_id, night, 'Hypnogram')
        if psg_file is None or hypno_file is None:
            print(f"Skipping subject {subject_id}, night {night}: Missing files")
            return None, None
        raw = mne.io.read_raw_edf(psg_file, preload=False, verbose=False)
        available_channels = [ch for ch in TARGET_CHANNELS if ch in raw.ch_names]
        if not available_channels:
            print(f"No target channels for subject {subject_id}, night {night}")
            return None, None
        raw.pick_channels(available_channels)
        raw.load_data()
        raw.filter(0.5, 40.0, l_trans_bandwidth=0.5, h_trans_bandwidth=10.0, verbose=False)
        raw.resample(SAMPLING_RATE, npad="auto")
        events = mne.make_fixed_length_events(raw, id=1, duration=EPOCH_DURATION)
        epochs_mne = mne.Epochs(raw, events, tmin=0, tmax=EPOCH_DURATION-1/raw.info['sfreq'],
                                picks=available_channels, baseline=None, preload=True)
        data = epochs_mne.get_data(units='uV')
        annotations = mne.read_annotations(hypno_file)
        labels = np.zeros(len(epochs_mne), dtype=int)
        stage_map = {'Sleep stage W': 0, 'Sleep stage 1': 1, 'Sleep stage 2': 2, 'Sleep stage 3': 3, 'Sleep stage 4': 3, 'Sleep stage R': 4}
        for annot in annotations:
            onset = int(annot['onset'] / EPOCH_DURATION)
            duration = int(annot['duration'] / EPOCH_DURATION)
            stage = annot['description']
            if stage in stage_map:
                for i in range(max(0, onset), min(len(epochs_mne), onset + duration)):
                    labels[i] = stage_map[stage]
        data = (data - np.mean(data, axis=(1, 2), keepdims=True)) / np.std(data, axis=(1, 2), keepdims=True)
        X = data.transpose(0, 2, 1)
        del raw, epochs_mne, data
        gc.collect()
        print(f"Processed subject {subject_id}, night {night}: {X.shape[0]} epochs")
        return X, labels
    except Exception as e:
        print(f"Error processing subject {subject_id}, night {night}: {e}")
        return None, None

class DataGenerator(Sequence):
    def __init__(self, X, y, batch_size, augment=True, class_weights=None):
        self.X = X.astype(np.float32)
        self.y = y.astype(np.int32)
        self.batch_size = batch_size
        self.augment = augment
        self.class_weights = class_weights
    def __len__(self):
        return int(np.ceil(len(self.X) / self.batch_size))
    def __getitem__(self, idx):
        start = idx * self.batch_size
        end = min(start + self.batch_size, len(self.X))
        X_batch = self.X[start:end]
        y_batch = self.y[start:end]
        if self.augment:
            X_batch = augment_data(X_batch).astype(np.float32)
        sample_weights = np.ones_like(y_batch, dtype=np.float32)
        if self.class_weights:
            sample_weights = np.array([self.class_weights[label] for label in y_batch], dtype=np.float32)
        return X_batch, y_batch, sample_weights


class TemporalAttention(layers.Layer):
    def __init__(self, heads=2, key_dim=16, **kwargs):
        super().__init__(**kwargs)
        self.multi_head = layers.MultiHeadAttention(num_heads=heads, key_dim=key_dim)
        self.norm = layers.LayerNormalization()
        self.add = layers.Add()
    def call(self, inputs):
        attn_output = self.multi_head(inputs, inputs)
        out = self.add([inputs, attn_output])
        return self.norm(out)


def build_lstm_attention_model(input_shape, nb_classes=5):
    inputs = layers.Input(shape=input_shape)
    x = layers.LSTM(128, return_sequences=True)(inputs)
    x = layers.DepthwiseConv1D(input_shape[1], depth_multiplier=2, padding='same', use_bias=False, depthwise_constraint=constraints.max_norm(1.0))(x)
    x = layers.BatchNormalization()(x)
    x = layers.Activation('elu')(x)
    x = layers.MaxPooling1D(pool_size=4)(x)
    x = layers.Dropout(0.25)(x)
    x = layers.SeparableConv1D(64, 16, padding='same', use_bias=False, depthwise_constraint=constraints.max_norm(1.0))(x)
    x = layers.BatchNormalization()(x)
    x = layers.Activation('elu')(x)
    x = layers.MaxPooling1D(pool_size=4)(x)
    x = TemporalAttention(name='temporal_attention') (x)
    x = layers.Dropout(0.25)(x)
    x = layers.GlobalAveragePooling1D()(x)
    x = layers.Dense(128, activation='relu')(x)
    x = layers.Dropout(0.3)(x)
    outputs = layers.Dense(nb_classes, activation='softmax', dtype='float32')(x)
    model = models.Model(inputs=inputs, outputs=outputs)
    model.compile(optimizer=tf.keras.optimizers.Adam(0.0005), loss='sparse_categorical_crossentropy', metrics=['accuracy'])
    return model

def build_eegnet_attention_model(input_shape, nb_classes=5, F1=8, D=2, F2=16, dropout_rate=0.25):
    inputs = layers.Input(shape=input_shape)
    x = layers.Conv1D(F1, 64, padding='same', use_bias=False, kernel_constraint=constraints.max_norm(1.0))(inputs)
    x = layers.BatchNormalization()(x)
    x = layers.DepthwiseConv1D(input_shape[1], depth_multiplier=D, padding='same', use_bias=False, depthwise_constraint=constraints.max_norm(1.0))(x)
    x = layers.BatchNormalization()(x)
    x = layers.Activation('elu')(x)
    x = layers.MaxPooling1D(pool_size=4)(x)
    x = layers.Dropout(dropout_rate)(x)
    x = layers.SeparableConv1D(F2, 16, padding='same', use_bias=False, depthwise_constraint=constraints.max_norm(1.0))(x)
    x = layers.BatchNormalization()(x)
    x = layers.Activation('elu')(x)
    x = layers.MaxPooling1D(pool_size=4)(x)
    x = TemporalAttention(name='temporal_attention') (x)
    x = layers.Dropout(dropout_rate)(x)
    x = layers.GlobalAveragePooling1D()(x)
    x = layers.Dense(nb_classes, activation='softmax', dtype='float32', kernel_constraint=constraints.max_norm(0.5))(x)
    model = models.Model(inputs=inputs, outputs=x)
    model.compile(optimizer=tf.keras.optimizers.Adam(0.0005), loss='sparse_categorical_crossentropy', metrics=['accuracy'])
    return model

def batched_predict(model, X, batch_size=BATCH_SIZE, training=False):
    preds = []
    for i in range(0, len(X), batch_size):
        batch = X[i:i+batch_size]
        pred = model(batch, training=training)
        preds.append(pred.numpy())
    return np.concatenate(preds, axis=0)

def build_mc_dropout_model(base_builder, input_shape, nb_classes=5, dropout_rate=0.25):
    model = base_builder(input_shape, nb_classes)
    return model

def predict_with_mc_dropout(model, X, n_samples=20, batch_size=BATCH_SIZE):
    preds = []
    for _ in range(n_samples):
        pred = batched_predict(model, X, batch_size, training=True)
        preds.append(pred)
    preds = np.stack(preds, axis=0)
    mean_pred = np.mean(preds, axis=0)
    uncertainty = np.std(preds, axis=0)
    return mean_pred, uncertainty

def train_deep_ensembles(base_builder, input_shape, nb_classes, train_gen, val_gen, n_ensembles=5):
    ensembles = []
    base_seed = 42
    early_stopping = EarlyStopping(monitor='val_loss', patience=3, restore_best_weights=True)
    for i in range(n_ensembles):
        seed = base_seed + i
        np.random.seed(seed)
        tf.random.set_seed(seed)
        random.seed(seed)
        os.environ['PYTHONHASHSEED'] = str(seed)
        model = base_builder(input_shape, nb_classes)
        model.fit(train_gen, validation_data=val_gen, epochs=EPOCHS, verbose=1, callbacks=[early_stopping])
        ensembles.append(model)
    return ensembles

def predict_with_deep_ensembles(ensembles, X, batch_size=BATCH_SIZE):
    preds = []
    for model in ensembles:
        pred = batched_predict(model, X, batch_size)
        preds.append(pred)
    preds = np.stack(preds, axis=0)
    mean_pred = np.mean(preds, axis=0)
    uncertainty = np.std(preds, axis=0)
    return mean_pred, uncertainty

def negative_log_likelihood(y_true, y_pred):
    y_true_onehot = tf.keras.utils.to_categorical(y_true, num_classes=y_pred.shape[1])
    return -np.mean(np.sum(y_true_onehot * np.log(y_pred + 1e-10), axis=1))

def brier_score(y_true, y_pred):
    y_true_onehot = tf.keras.utils.to_categorical(y_true, num_classes=y_pred.shape[1])
    return np.mean(np.sum((y_pred - y_true_onehot)**2, axis=1))

def expected_calibration_error(y_true, y_pred, num_bins=15):
    pred_classes = np.argmax(y_pred, axis=1)
    prob_max = np.max(y_pred, axis=1)
    correct = (pred_classes == y_true).astype(np.float32)

    bins = np.linspace(0, 1, num_bins + 1)
    indices = np.digitize(prob_max, bins, right=True)

    ece = 0
    for b in range(1, num_bins + 1):
        mask = (indices == b)
        if np.any(mask):
            acc = np.mean(correct[mask])
            conf = np.mean(prob_max[mask])
            ece += np.abs(acc - conf) * np.sum(mask) / len(y_true)
    return ece

def plot_training_curves(history, model_name):
    plt.figure(figsize=(12, 4))
    plt.subplot(1, 2, 1)
    plt.plot(history.history['accuracy'], label='Train Accuracy')
    plt.plot(history.history['val_accuracy'], label='Validation Accuracy')
    plt.title(f'{model_name} - Model Accuracy')
    plt.xlabel('Epoch')
    plt.ylabel('Accuracy')
    plt.legend()
    plt.subplot(1, 2, 2)
    plt.plot(history.history['loss'], label='Train Loss')
    plt.plot(history.history['val_loss'], label='Validation Loss')
    plt.title(f'{model_name} - Model Loss')
    plt.xlabel('Epoch')
    plt.ylabel('Loss')
    plt.legend()
    plt.tight_layout()
    plt.savefig(f'training_curves_{model_name}.png')
    plt.close()

def plot_calibration_curve(y_true, y_pred, model_name, num_bins=15):
    prob_max = np.max(y_pred, axis=1)
    bins = np.linspace(0, 1, num_bins + 1)
    bin_centers = (bins[1:] + bins[:-1]) / 2
    accs = []
    confs = []
    bin_centers_filled = []
    for i in range(num_bins):
        mask = (prob_max >= bins[i]) & (prob_max < bins[i+1])
        if np.sum(mask) > 0:
            acc = np.mean(y_true[mask] == np.argmax(y_pred[mask], axis=1))
            conf = np.mean(prob_max[mask])
            accs.append(acc)
            confs.append(conf)
            bin_centers_filled.append(bin_centers[i])
    plt.figure(figsize=(8, 6))
    plt.plot([0,1], [0,1], 'k--', label='Perfect Calibration')
    plt.bar(bin_centers_filled, accs, width=(1/num_bins), alpha=0.5, label='Accuracy')
    plt.plot(bin_centers_filled, confs, 'r-', label='Confidence')
    plt.xlabel('Confidence')
    plt.ylabel('Accuracy')
    plt.title(f'{model_name} - Calibration Curve')
    plt.legend()
    plt.savefig(f'calibration_curve_{model_name}.png')
    plt.close()

def plot_uncertainty_heatmap(y_true, uncertainty, model_name):
    cm_unc = np.zeros((nb_classes, nb_classes))
    for i in range(nb_classes):
        mask = y_true == i
        if np.sum(mask) > 0:
            avg_unc = np.mean(uncertainty[mask], axis=0)
            cm_unc[i, :] = avg_unc
    stage_names = ['Wake', 'N1', 'N2', 'N3', 'REM']
    plt.figure(figsize=(8, 6))
    sns.heatmap(cm_unc, annot=True, fmt='.4f', cmap='viridis', xticklabels=stage_names, yticklabels=stage_names)
    plt.xlabel('Class')
    plt.ylabel('True Stage')
    plt.title(f'{model_name} - Average Uncertainty per Class')
    plt.savefig(f'uncertainty_heatmap_{model_name}.png')
    plt.close()

def plot_uncertainty_distribution(y_true, mean_pred, model_name):
    ent = -np.sum(mean_pred * np.log(mean_pred + 1e-10), axis=1)
    data = []
    for i in range(nb_classes):
        mask = y_true == i
        if np.sum(mask) > 0:
            data.append(ent[mask])
        else:
            data.append([])
    stage_names = ['Wake', 'N1', 'N2', 'N3', 'REM']
    plt.figure(figsize=(10, 6))
    sns.violinplot(data=data)
    plt.xticks(range(nb_classes), stage_names)
    plt.xlabel('Sleep Stage')
    plt.ylabel('Predictive Entropy')
    plt.title(f'{model_name} - Uncertainty Distribution')
    plt.savefig(f'uncertainty_violin_{model_name}.png')
    plt.close()

def plot_attention_weights(model, X_test, y_test, model_name, num_samples=5):
    stage_names = ['Wake', 'N1', 'N2', 'N3', 'REM']
    try:
        intermediate_model = models.Model(inputs=model.inputs, outputs=model.get_layer('temporal_attention').input)
        multi_head = model.get_layer('temporal_attention').multi_head
    except ValueError:
        print(f"No temporal_attention layer found for {model_name}")
        return
    selected_indices = []
    for i in range(nb_classes):
        mask = np.where(y_test == i)[0]
        if len(mask) > 0:
            selected_indices.append(np.random.choice(mask))
    if not selected_indices:
        return
    X_samples = X_test[selected_indices]
    attn_inputs = intermediate_model([X_samples])
    _, attn_scores = multi_head(attn_inputs, attn_inputs, return_attention_scores=True)
    for idx, sample_idx in enumerate(selected_indices):
        stage = stage_names[y_test[sample_idx]]
        plt.figure(figsize=(10, 8))
        for head in range(attn_scores.shape[1]):
            plt.subplot(1, attn_scores.shape[1], head + 1)
            sns.heatmap(attn_scores[idx, head], cmap='viridis')
            plt.title(f'Head {head + 1}')
        plt.suptitle(f'{model_name} - Attention Weights for {stage} Sample')
        plt.tight_layout()
        plt.savefig(f'attention_weights_{model_name}_{stage}.png')
        plt.close()

def plot_brier_comparison(results):
    names = list(results.keys())
    briers = [met['brier'] for met in results.values()]
    data = pd.DataFrame({'Model': names, 'Brier Score': briers})
    plt.figure(figsize=(10, 6))
    sns.barplot(data=data, x='Model', y='Brier Score')
    plt.title('Brier Score Comparison Across Models')
    plt.xticks(rotation=45)
    plt.savefig('brier_scores_comparison.png')
    plt.close()

def evaluate_model(model_name, predict_func, X_test, y_test):
    mean_pred, uncertainty = predict_func(X_test)
    test_acc = np.mean(np.argmax(mean_pred, axis=1) == y_test)
    nll = negative_log_likelihood(y_test, mean_pred)
    brier = brier_score(y_test, mean_pred)
    ece = expected_calibration_error(y_test, mean_pred)

    print(f"\n{model_name} - Test Accuracy: {test_acc:.4f}")
    print(f"{model_name} - NLL: {nll:.4f}")
    print(f"{model_name} - Brier Score: {brier:.4f}")
    print(f"{model_name} - ECE: {ece:.4f}")

    y_pred_classes = np.argmax(mean_pred, axis=1)
    precision, recall, f1, _ = precision_recall_fscore_support(y_test, y_pred_classes, average=None)
    stage_names = ['Wake', 'N1', 'N2', 'N3', 'REM']
    print(f"\n{model_name} - Per-class Metrics:")
    for i, stage in enumerate(stage_names):
        print(f"{stage}: Precision={precision[i]:.4f}, Recall={recall[i]:.4f}, F1={f1[i]:.4f}")
    cm = confusion_matrix(y_test, y_pred_classes)
    plt.figure(figsize=(8, 6))
    sns.heatmap(cm, annot=True, fmt='d', cmap='Blues', xticklabels=stage_names, yticklabels=stage_names)
    plt.xlabel('Predicted')
    plt.ylabel('True')
    plt.title(f'{model_name} - Confusion Matrix')
    plt.savefig(f'confusion_matrix_{model_name}.png')
    plt.close()

    plot_calibration_curve(y_test, mean_pred, model_name)
    plot_uncertainty_heatmap(y_test, uncertainty, model_name)
    plot_uncertainty_distribution(y_test, mean_pred, model_name)

    return {'acc': test_acc, 'nll': nll, 'brier': brier, 'ece': ece}

def data_generator(available, batch_size=2000):
    for subject_id, night in available:
        X, y = process_subject_night(subject_id, night)
        if X is None or y is None:
            continue
        for i in range(0, len(X), batch_size):
            yield X[i:i+batch_size], y[i:i+batch_size]
        del X, y
        gc.collect()

def run_pipeline():
    available = get_available_subjects()
    if not available:
        return
    X_train, y_train, X_test, y_test = [], [], [], []
    for X_batch, y_batch in tqdm(data_generator(available), desc="Processing data"):
        if X_batch is None or y_batch is None:
            continue
        class_counts = np.bincount(y_batch)
        stratify = y_batch if min(class_counts[class_counts > 0]) >= 2 else None
        X_tr, X_te, y_tr, y_te = train_test_split(X_batch, y_batch, test_size=0.2, stratify=stratify, random_state=42)
        X_train.append(X_tr); y_train.append(y_tr)
        X_test.append(X_te); y_test.append(y_te)
        del X_batch, y_batch
        gc.collect()
    if not X_train:
        return
    X_train = np.concatenate(X_train)
    y_train = np.concatenate(y_train)
    X_test = np.concatenate(X_test)
    y_test = np.concatenate(y_test)
    class_weights = compute_class_weight('balanced', classes=np.unique(y_train), y=y_train)
    class_weight_dict = dict(enumerate(class_weights))
    train_generator = DataGenerator(X_train, y_train, BATCH_SIZE, augment=True, class_weights=class_weight_dict)
    val_generator = DataGenerator(X_test, y_test, BATCH_SIZE, augment=False, class_weights=class_weight_dict)

    input_shape = (X_train.shape[1], X_train.shape[2])

    base_builders = [('EEGNet', build_eegnet_attention_model), ('LSTM', build_lstm_attention_model)]
    uncertainty_methods = [
        ('MC_Dropout', build_mc_dropout_model, lambda model, X: predict_with_mc_dropout(model, X)),
        ('Deep_Ensembles', train_deep_ensembles, lambda ensembles, X: predict_with_deep_ensembles(ensembles, X))
    ]

    results = {}
    early_stopping = EarlyStopping(monitor='val_loss', patience=3, restore_best_weights=True)
    for base_name, base_builder in base_builders:
        for unc_name, build_func, predict_func in uncertainty_methods:
            model_name = f"{base_name}_{unc_name}"
            print(f"\nTraining {model_name}...")
            if unc_name == 'Deep_Ensembles':
                models = build_func(base_builder, input_shape, nb_classes, train_generator, val_generator)
                metrics = evaluate_model(model_name, lambda X: predict_func(models, X), X_test, y_test)
                plot_attention_weights(models[0], X_test, y_test, model_name)
            else:
                model = build_func(base_builder, input_shape, nb_classes)
                history = model.fit(train_generator, validation_data=val_generator, epochs=EPOCHS, verbose=1, callbacks=[early_stopping])
                plot_training_curves(history, model_name)
                metrics = evaluate_model(model_name, lambda X: predict_func(model, X), X_test, y_test)
                plot_attention_weights(model, X_test, y_test, model_name)
            results[model_name] = metrics

    print("\nModel Comparison for Uncertainty Estimation:")
    print("Sorted by ECE (lower is better):")
    for name, met in sorted(results.items(), key=lambda x: x[1]['ece']):
        print(f"{name}: Acc={met['acc']:.4f}, NLL={met['nll']:.4f}, Brier={met['brier']:.4f}, ECE={met['ece']:.4f}")

    plot_brier_comparison(results)

    best_model = min(results, key=lambda k: results[k]['ece'])
    print(f"\nBest fit model for uncertainty estimation: {best_model} with ECE {results[best_model]['ece']:.4f}")

if __name__ == "__main__":
    run_pipeline()
```
