## Supplementary figures and images for "Trustworthy Sleep Staging from EEG: Deep Ensembles, MC Dropout, and Predictive Calibration"

### attention_weights_EEGNet_Deep_Ensembles_N1.png

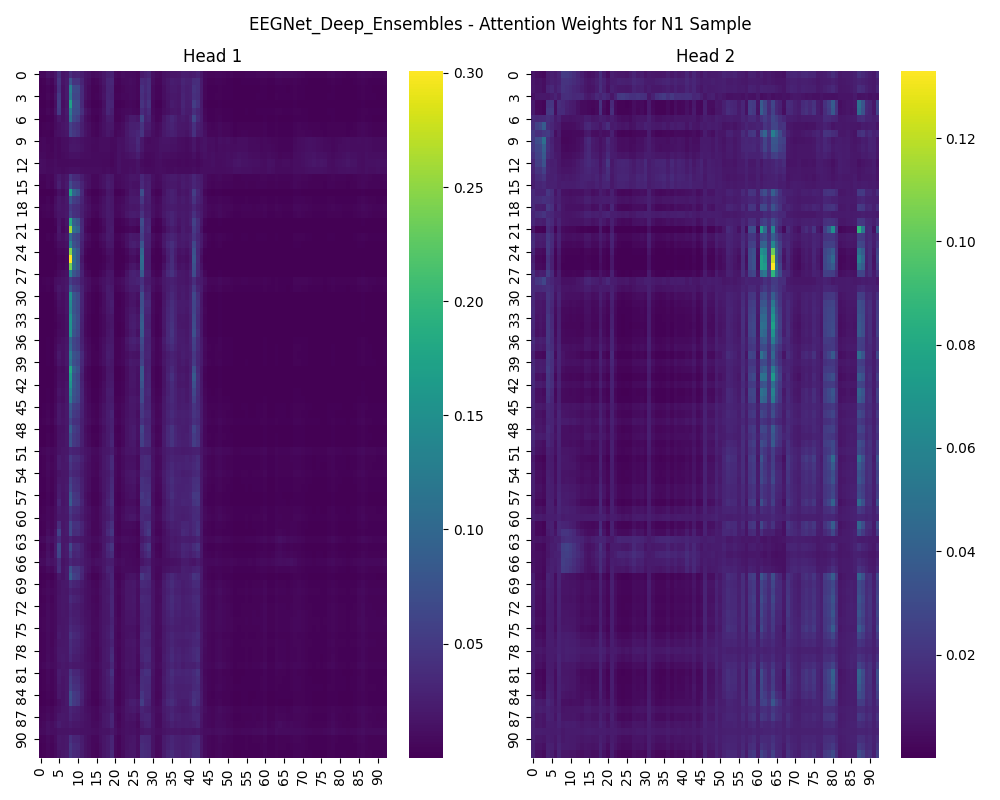

### attention_weights_EEGNet_Deep_Ensembles_N2.png

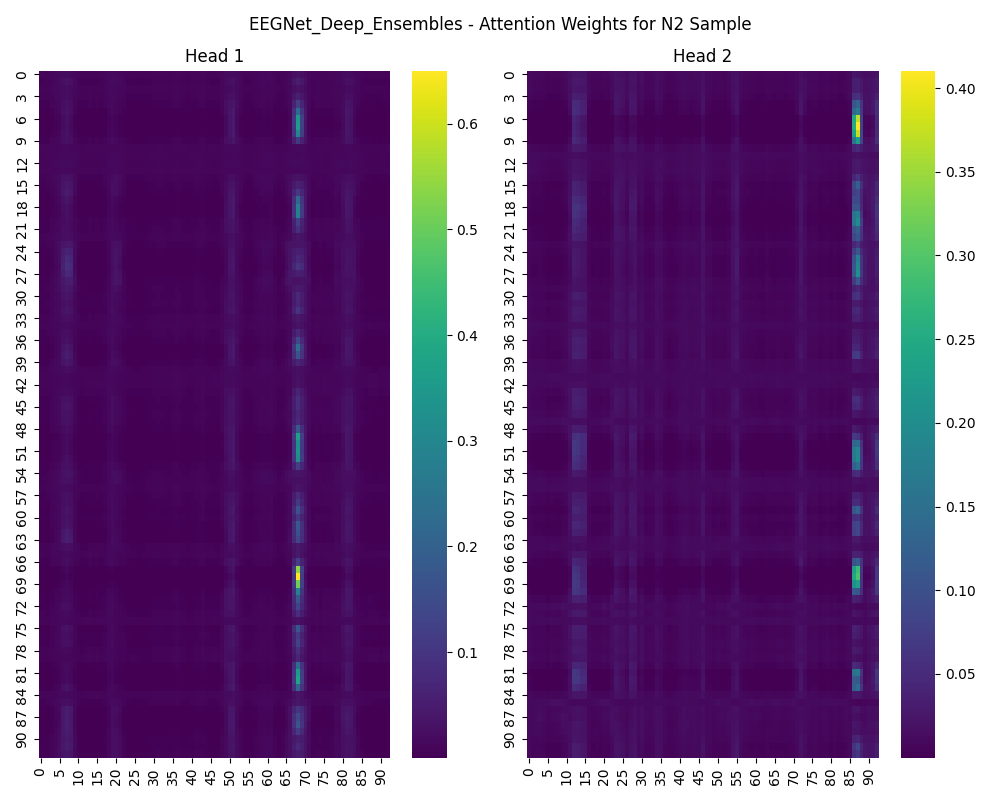

### attention_weights_EEGNet_Deep_Ensembles_N3.png

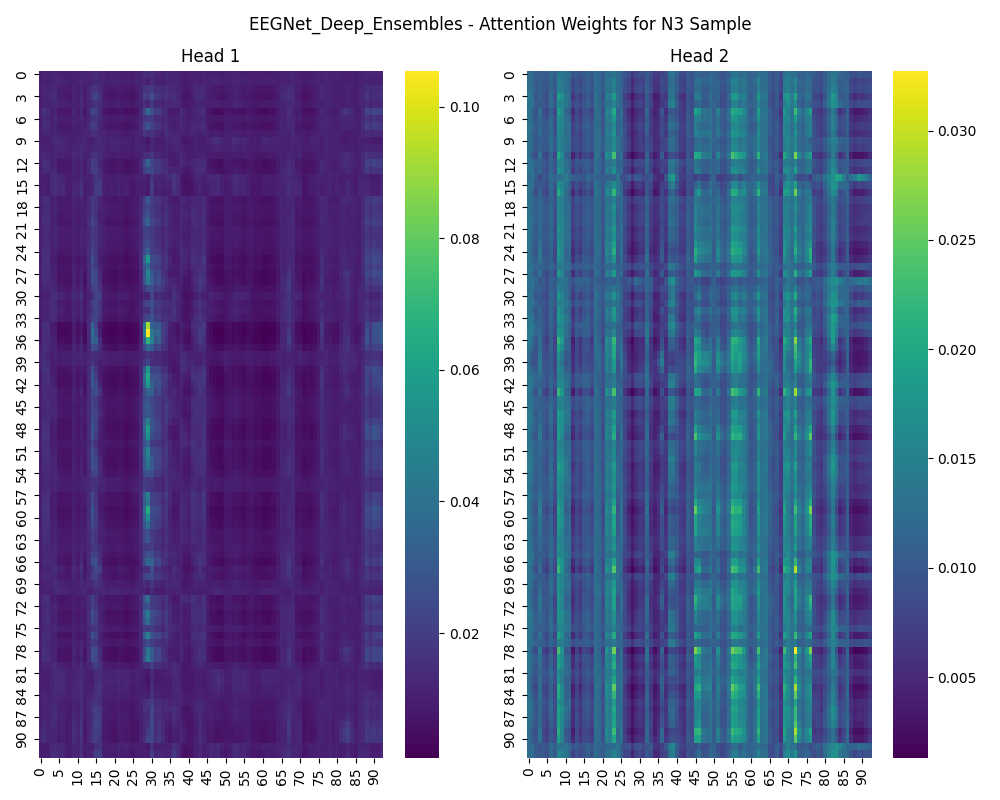

### attention_weights_EEGNet_Deep_Ensembles_REM.png

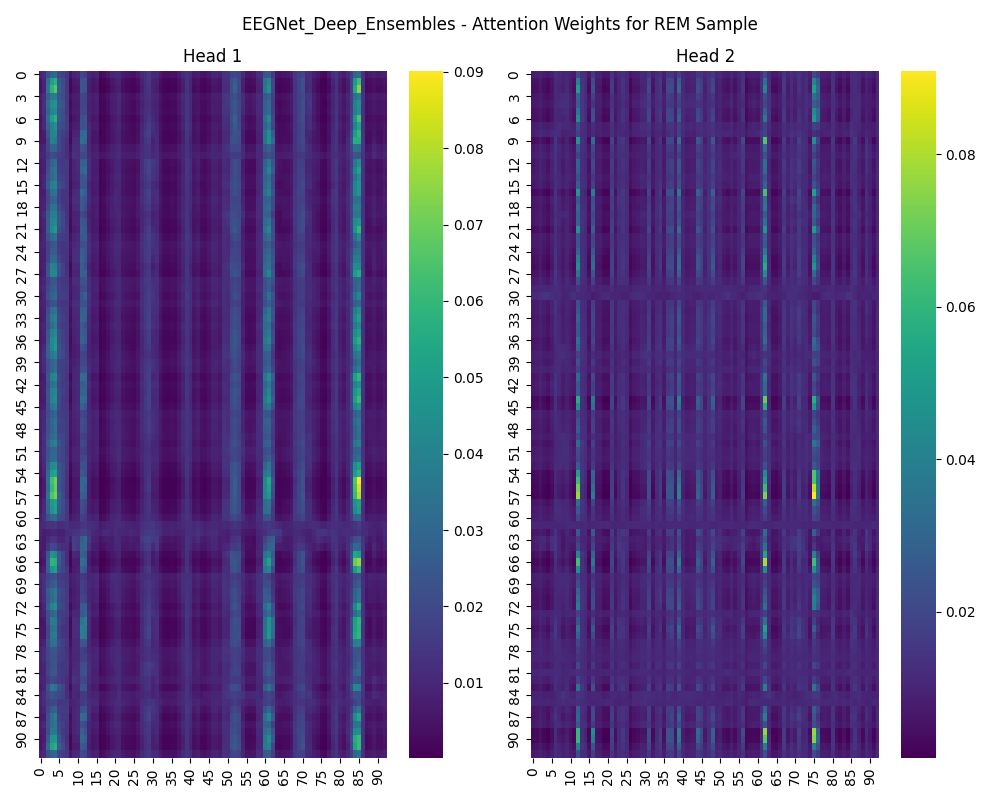

### attention_weights_EEGNet_Deep_Ensembles_Wake.png

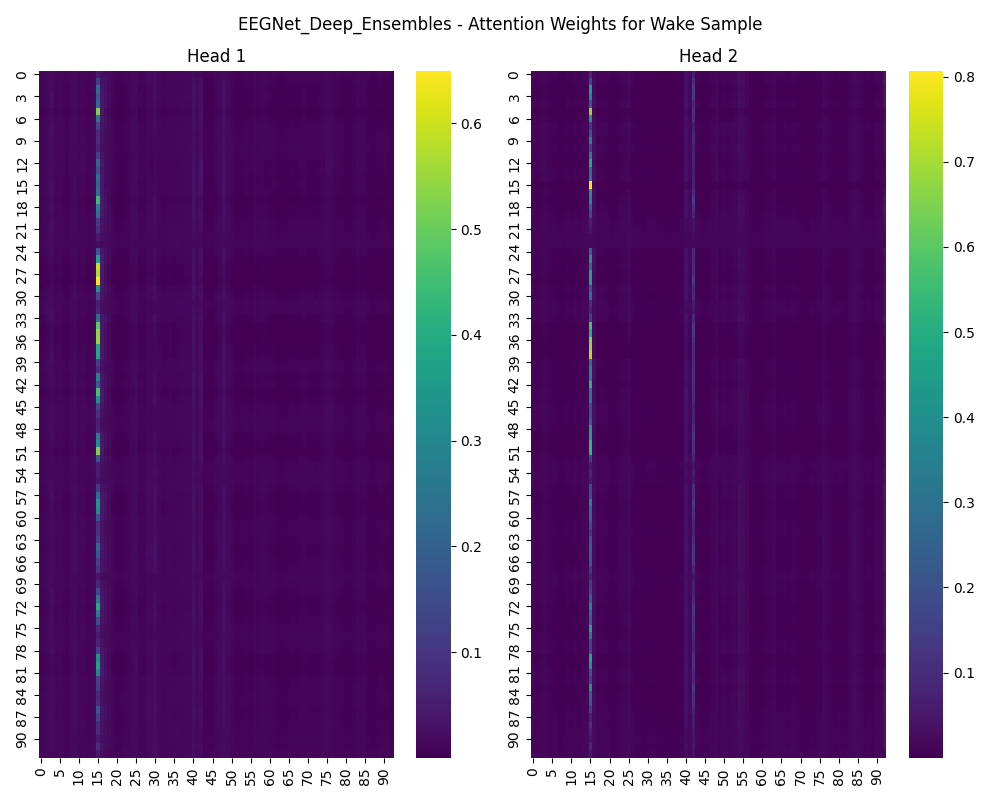

### attention_weights_EEGNet_MC_Dropout_N1.png

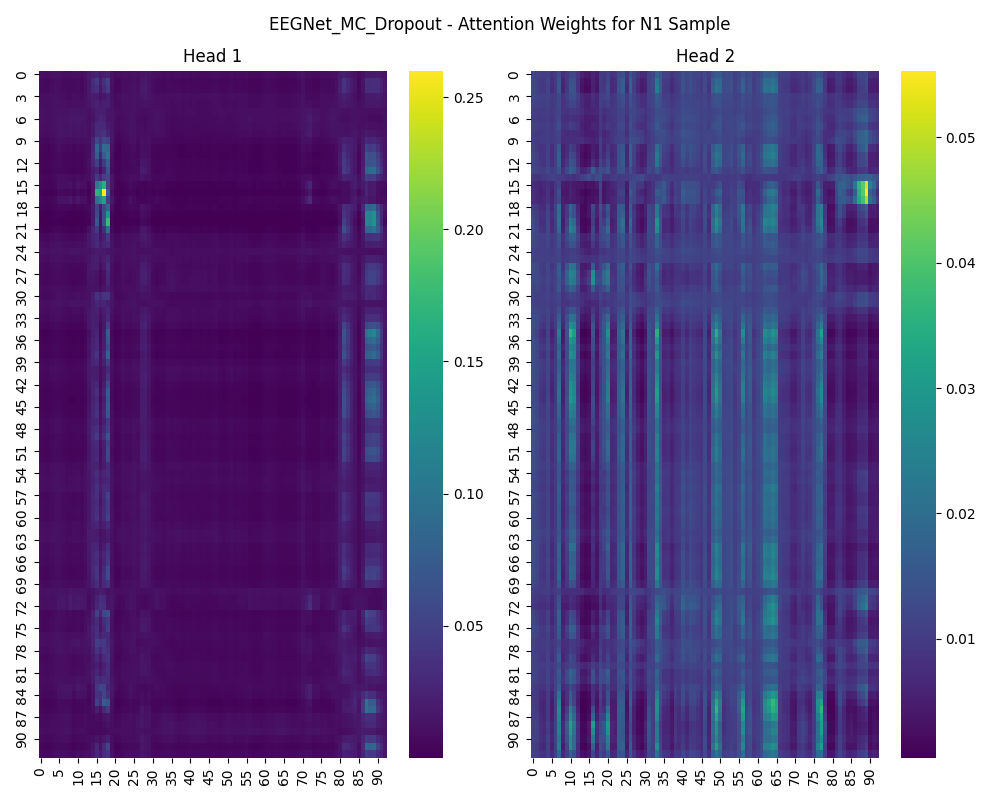

### attention_weights_EEGNet_MC_Dropout_N2.png

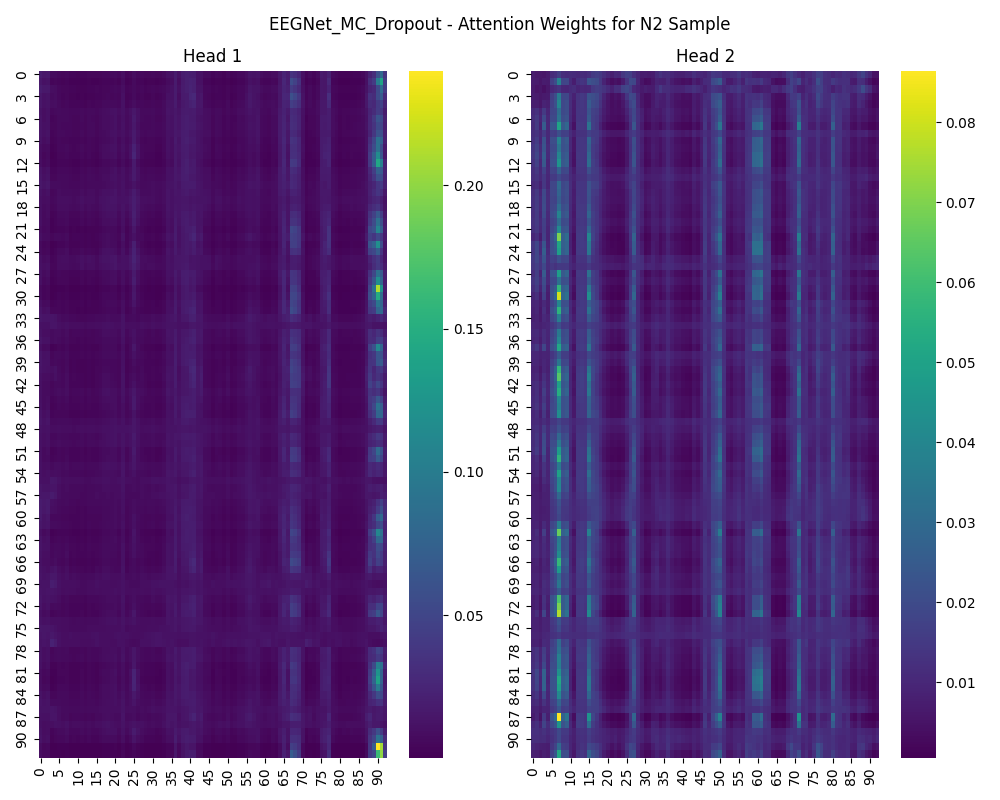

### attention_weights_EEGNet_MC_Dropout_N3.png

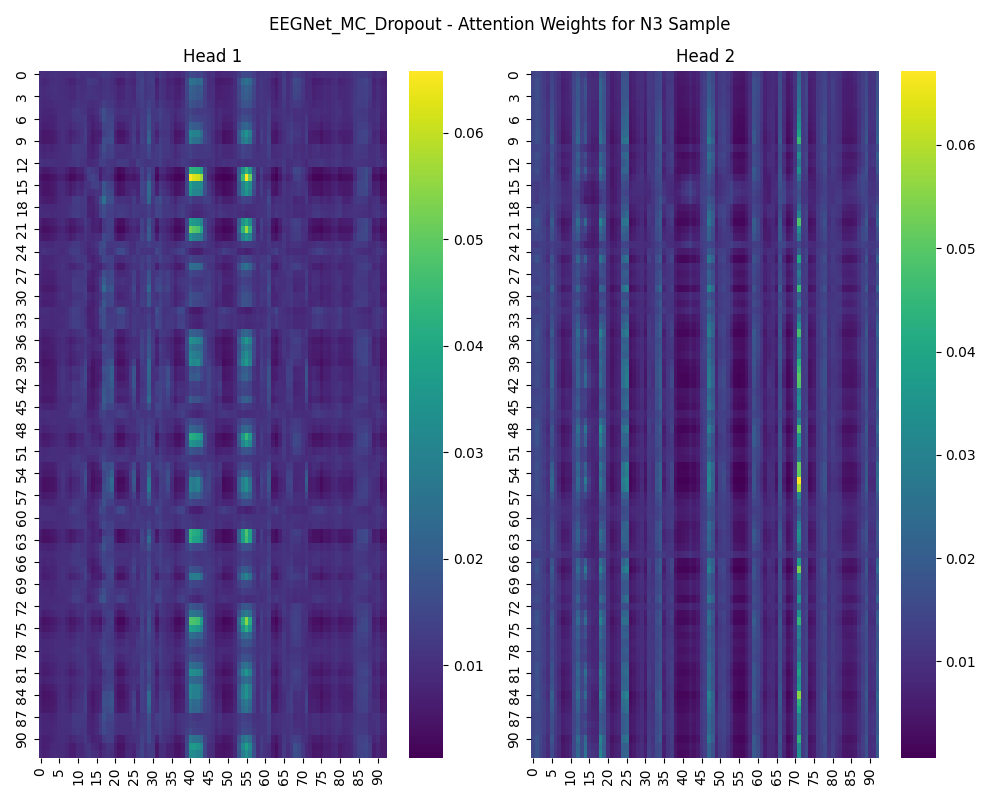

### attention_weights_EEGNet_MC_Dropout_REM.png

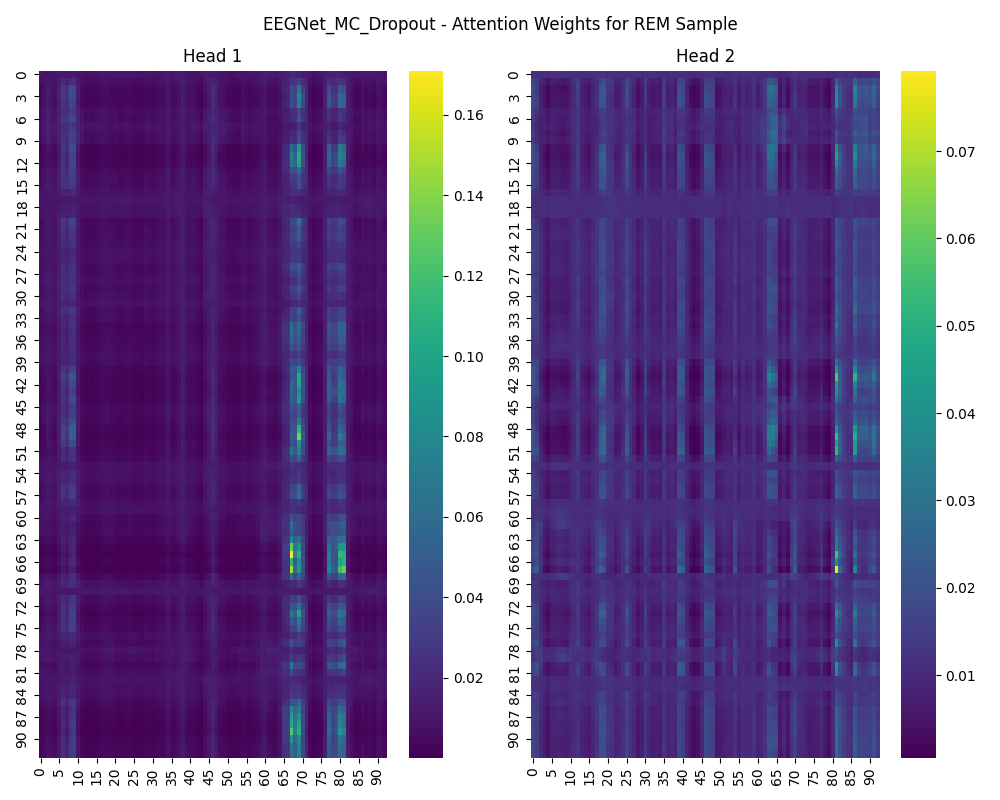

### attention_weights_EEGNet_MC_Dropout_Wake.png

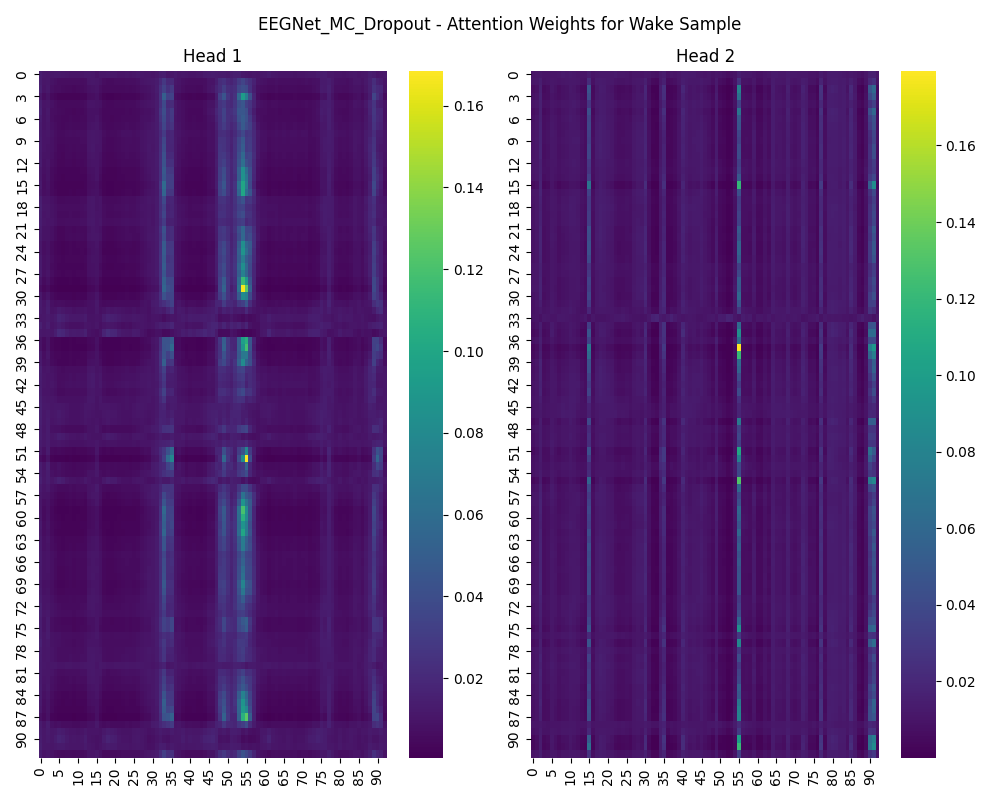

### attention_weights_LSTM_Deep_Ensembles_N1.png

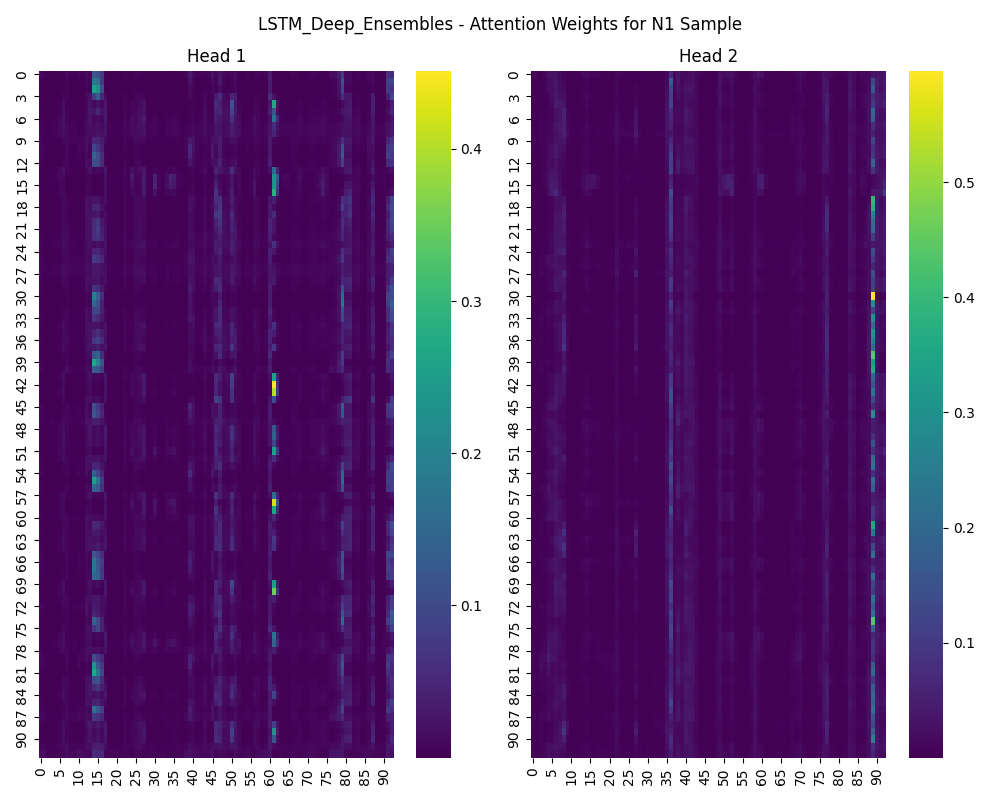

### attention_weights_LSTM_Deep_Ensembles_N2.png

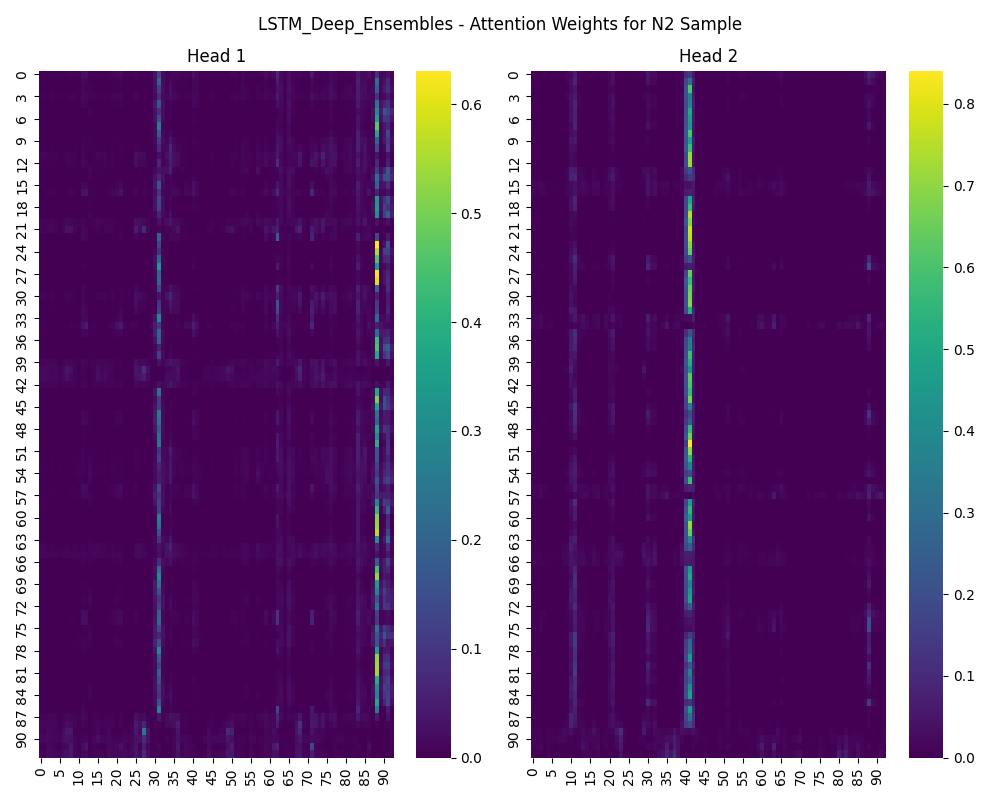

### attention_weights_LSTM_Deep_Ensembles_N3.png

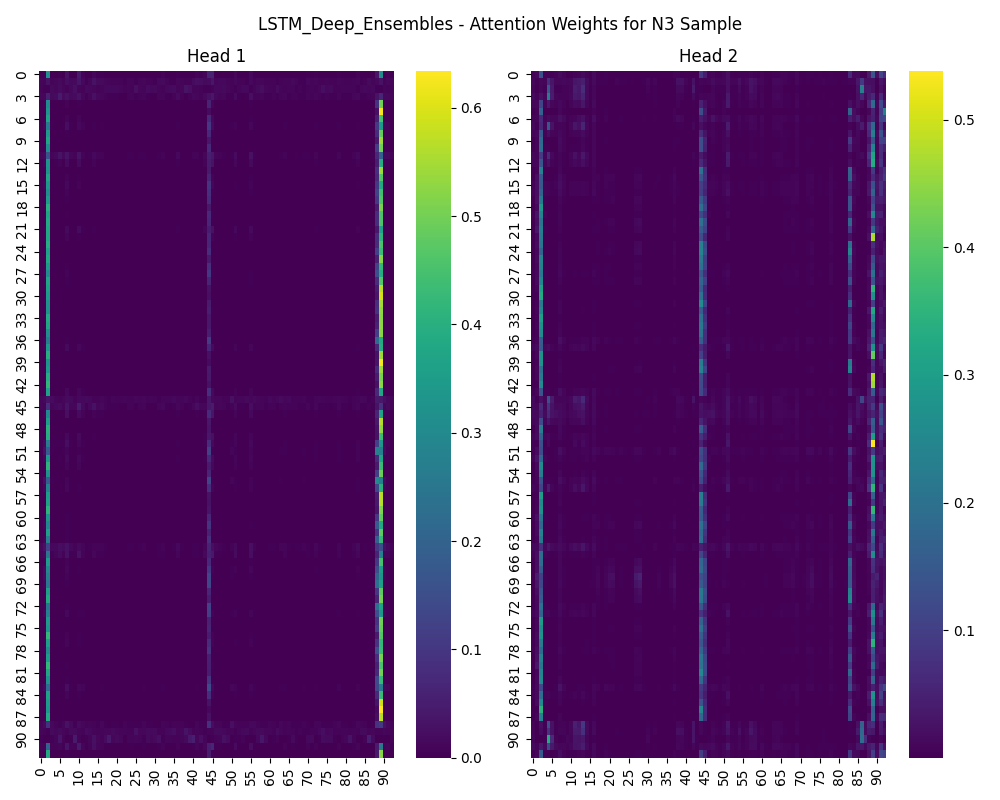

### attention_weights_LSTM_Deep_Ensembles_REM.png

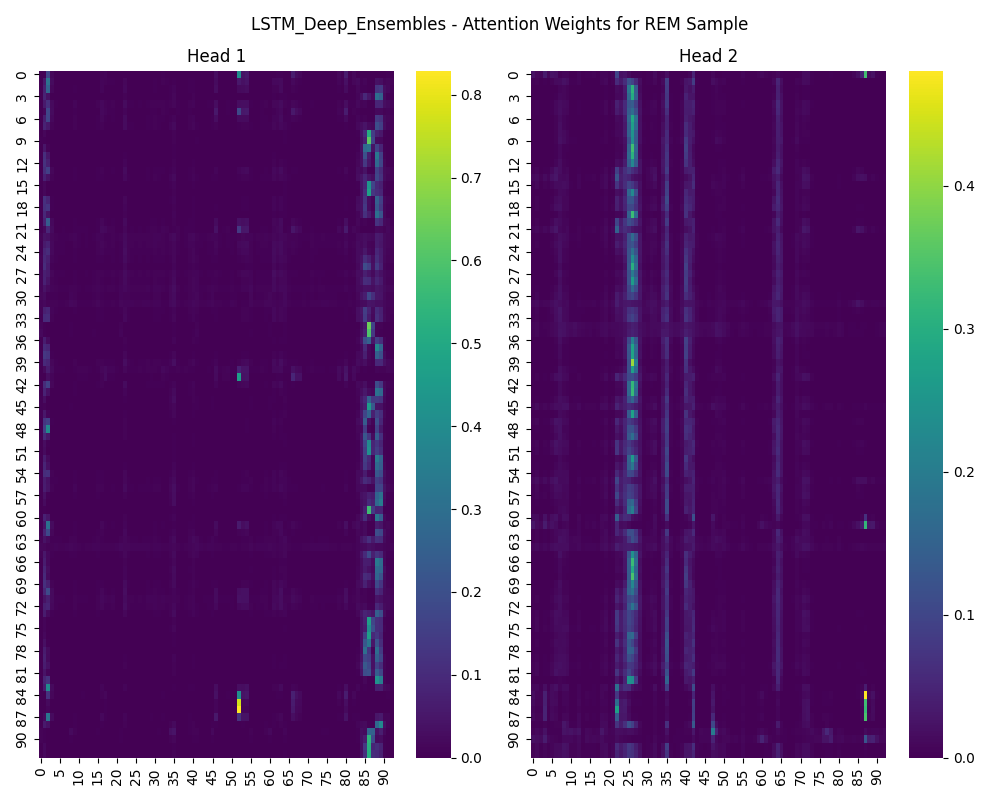

### attention_weights_LSTM_Deep_Ensembles_Wake.png

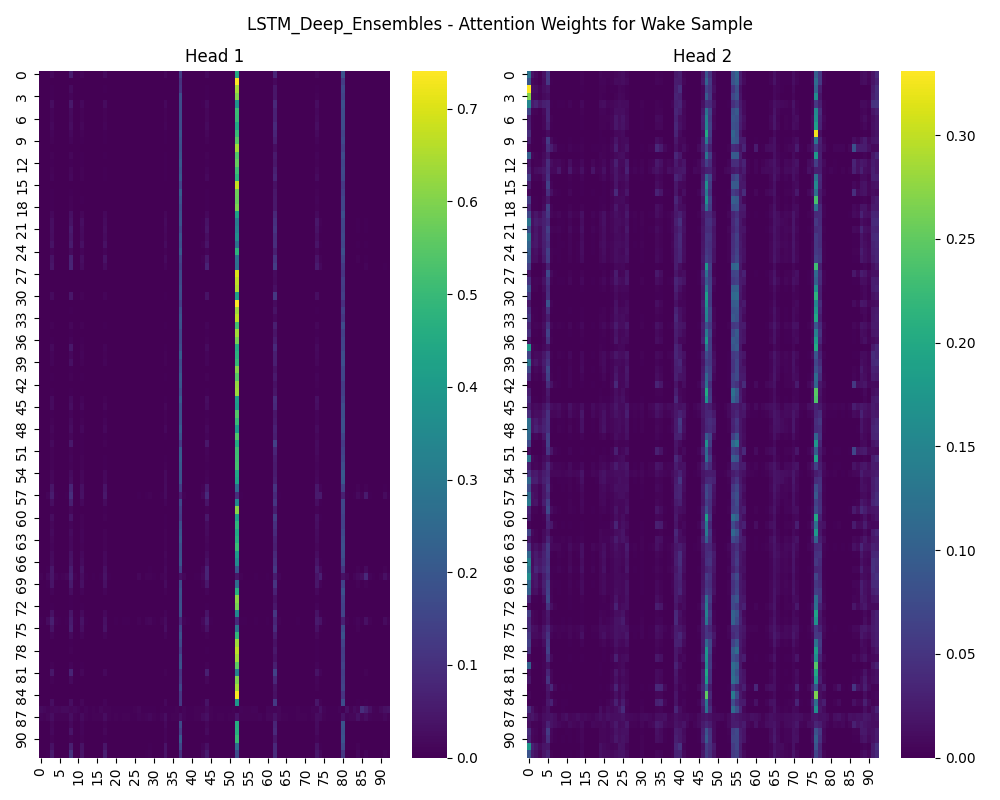

### attention_weights_LSTM_MC_Dropout_N1.png

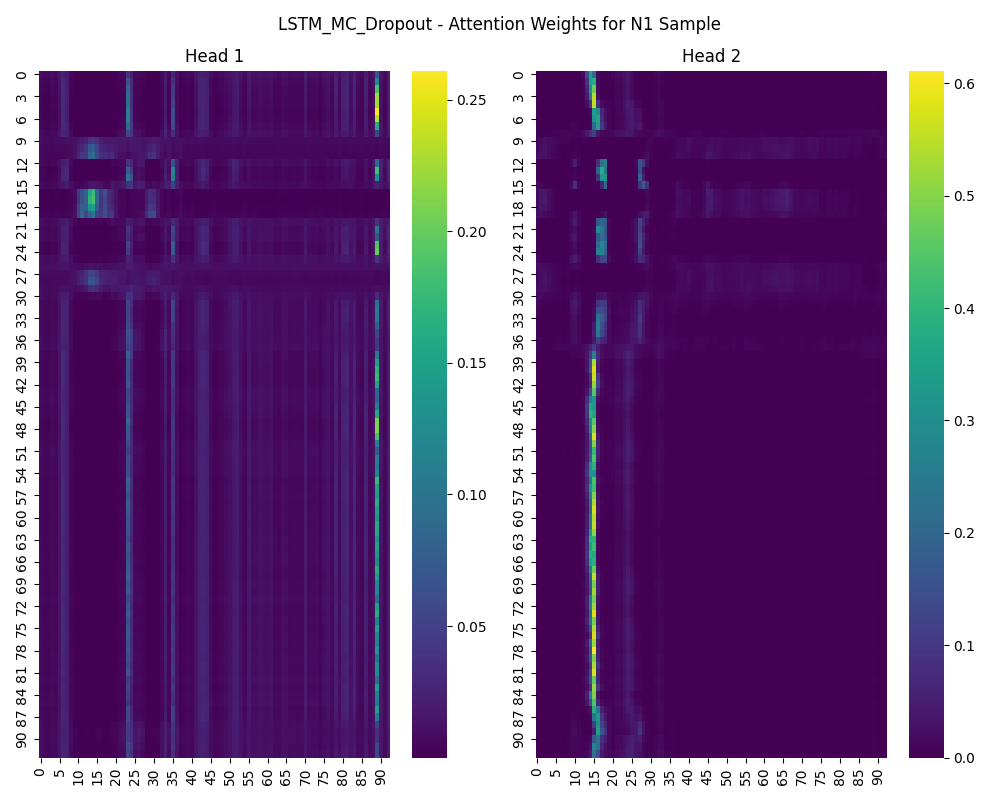

### attention_weights_LSTM_MC_Dropout_N2.png

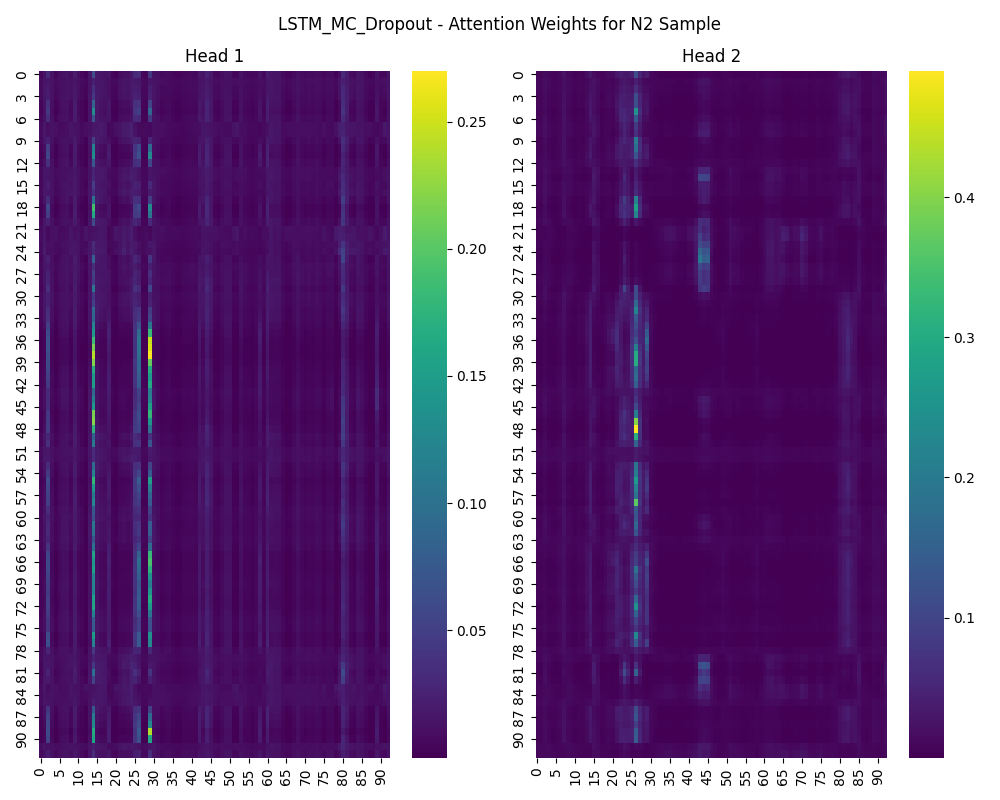

### attention_weights_LSTM_MC_Dropout_N3.png

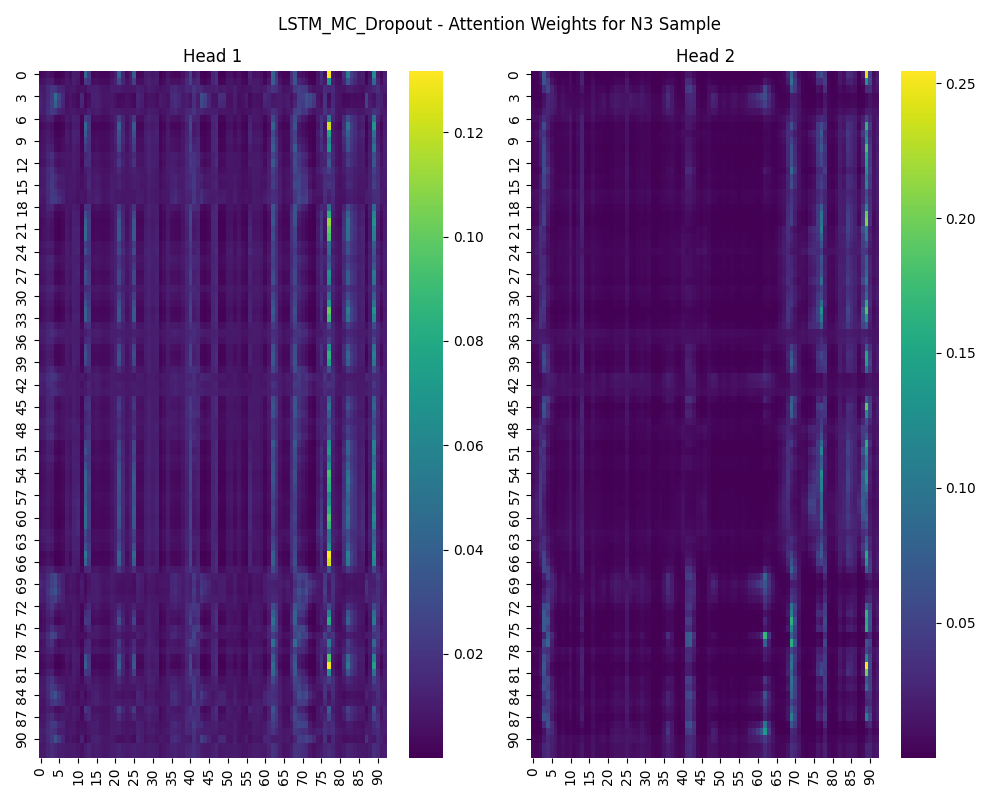

### attention_weights_LSTM_MC_Dropout_REM.png

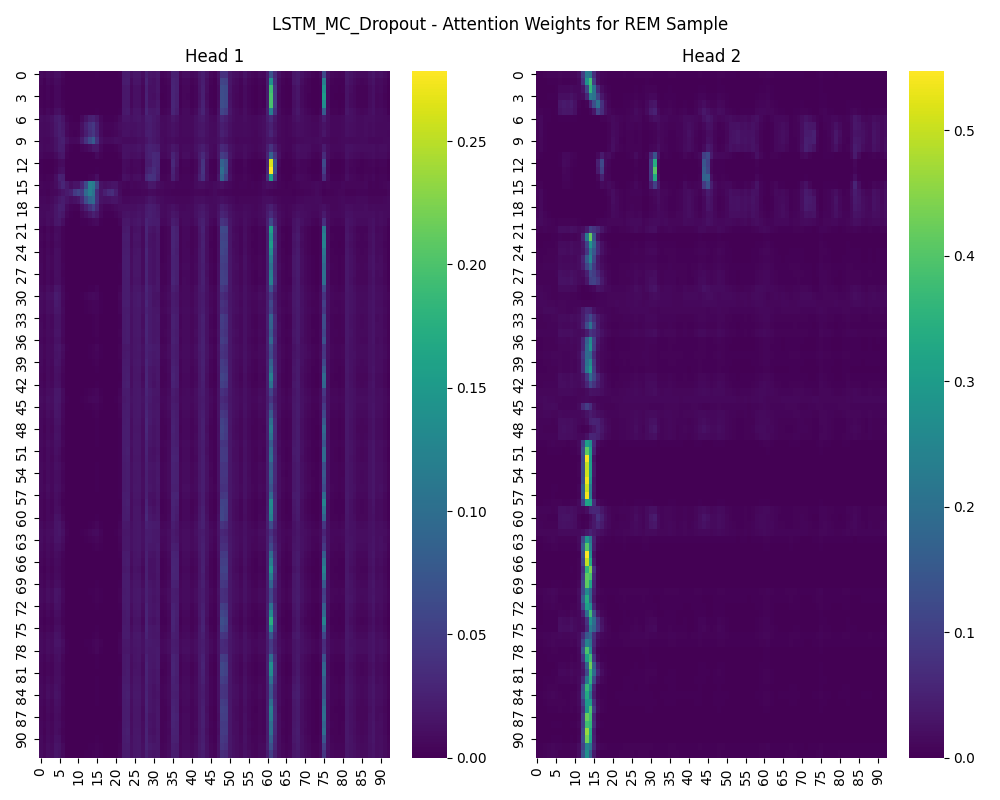

### attention_weights_LSTM_MC_Dropout_Wake.png

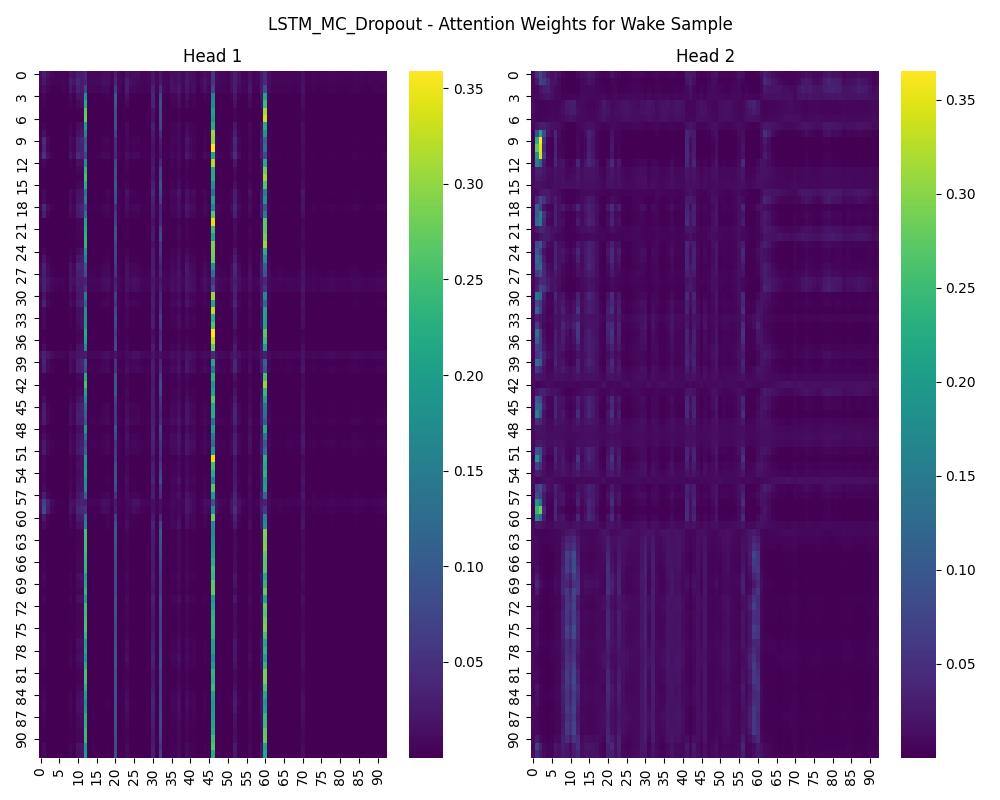

### brier_scores_comparison.png

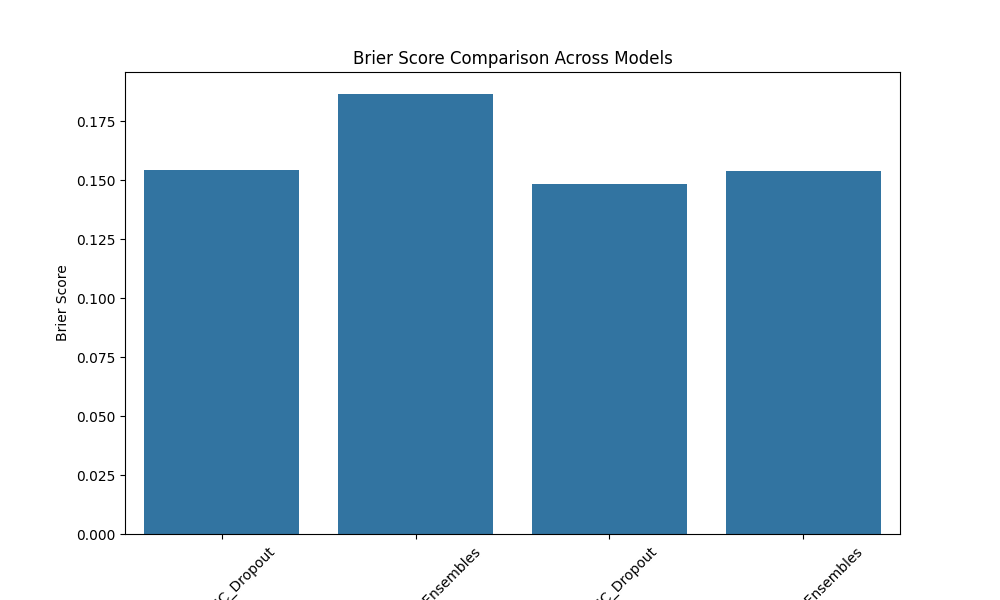

### calibration_curve_EEGNet_Deep_Ensembles.png

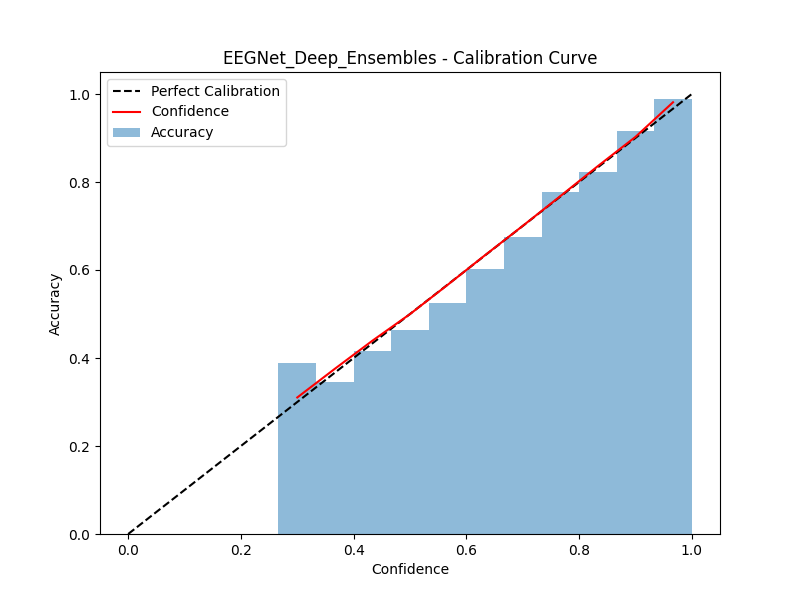

### calibration_curve_EEGNet_MC_Dropout.png

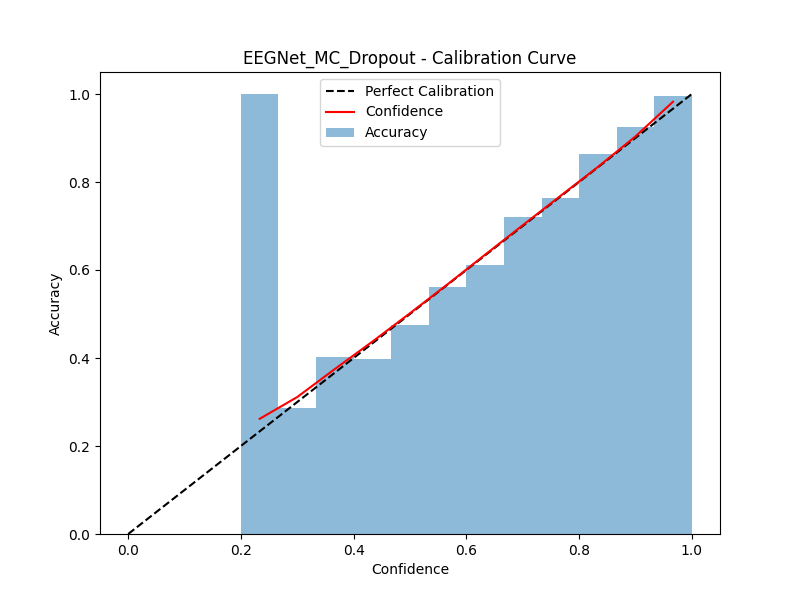

### calibration_curve_LSTM_Deep_Ensembles.png

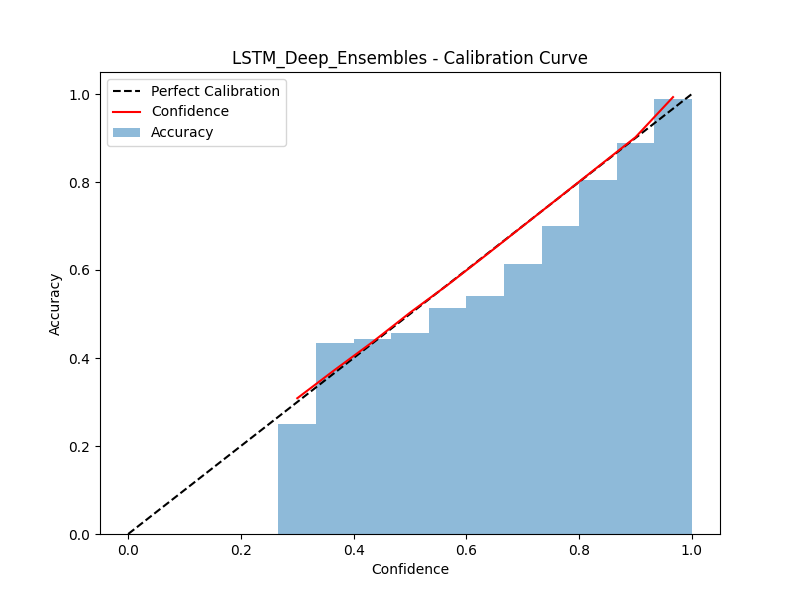

### calibration_curve_LSTM_MC_Dropout.png

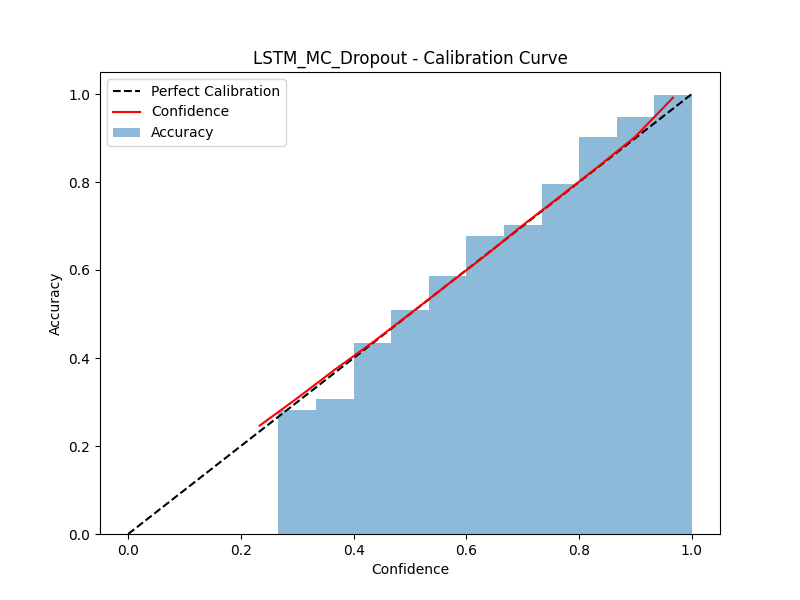

### confusion_matrix_EEGNet_Deep_Ensembles.png

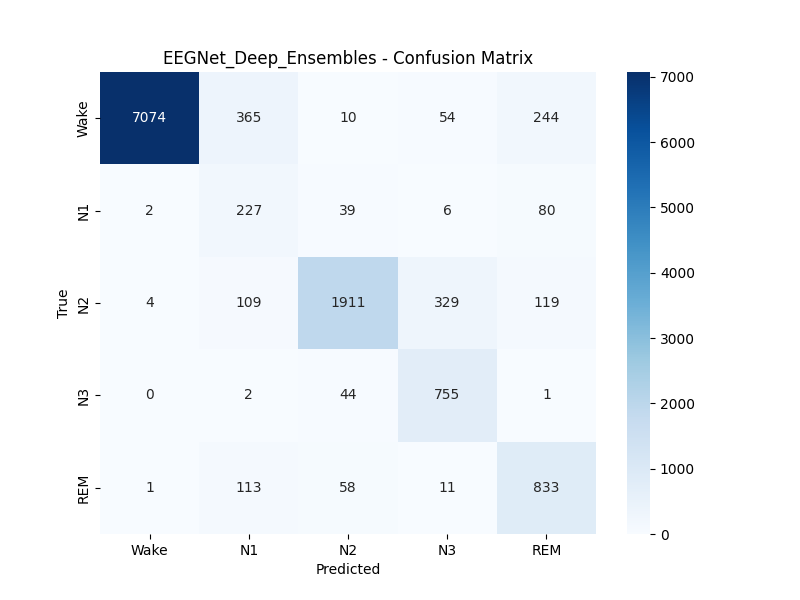

### confusion_matrix_EEGNet_MC_Dropout.png

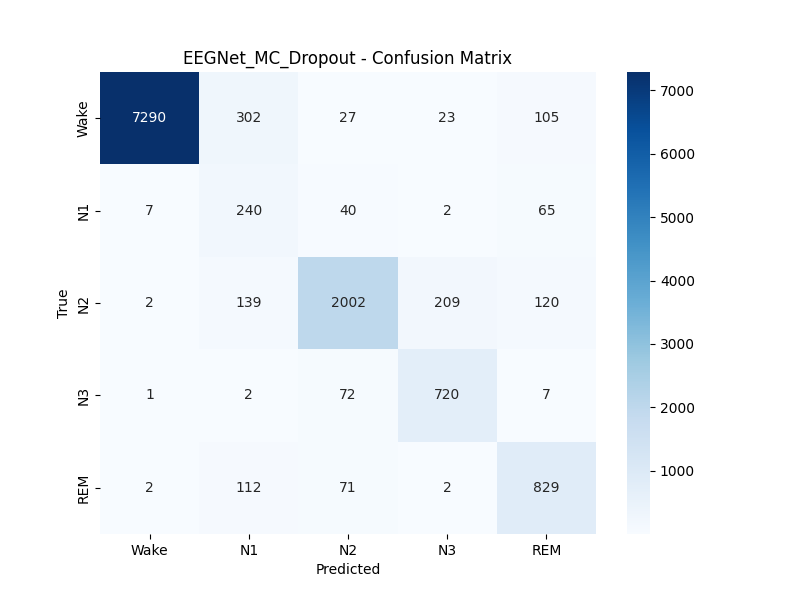

### confusion_matrix_LSTM_Deep_Ensembles.png

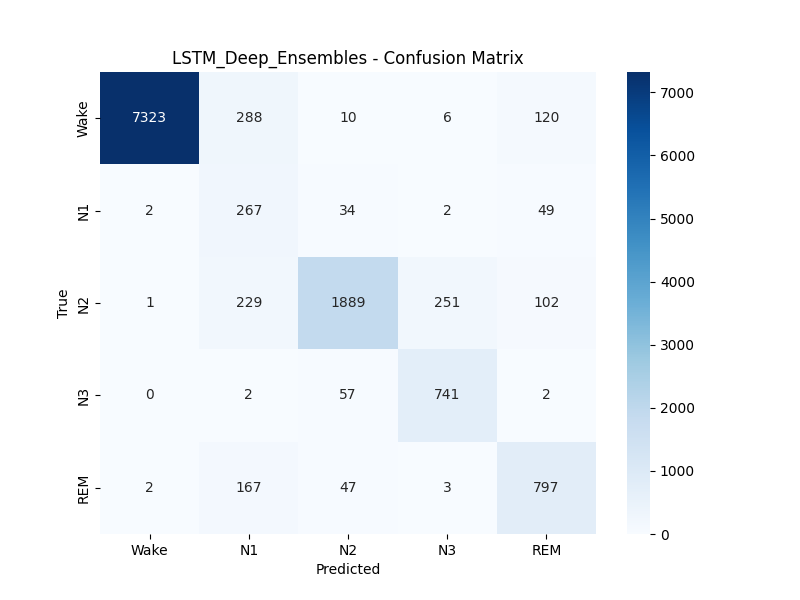

### confusion_matrix_LSTM_MC_Dropout.png

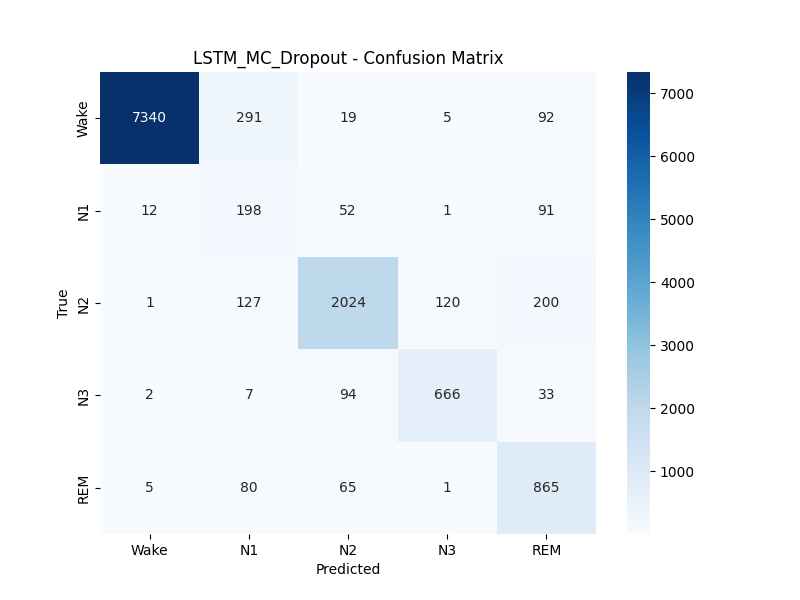

### training_curves_EEGNet_MC_Dropout.png

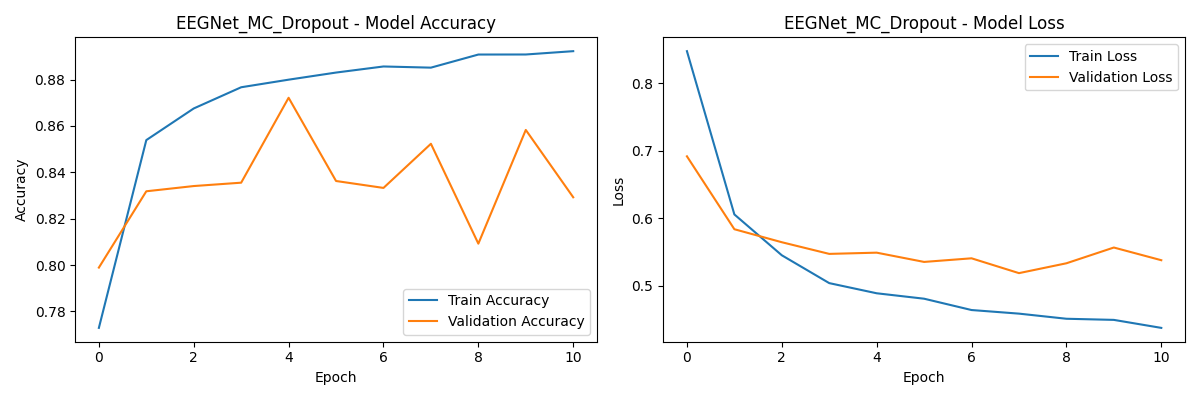

### training_curves_LSTM_MC_Dropout.png

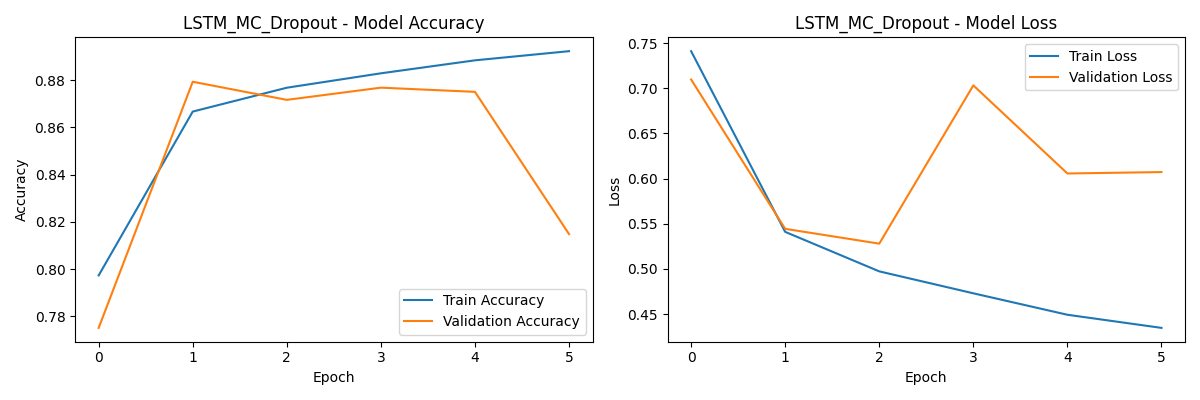

### uncertainty_heatmap_EEGNet_Deep_Ensembles.png

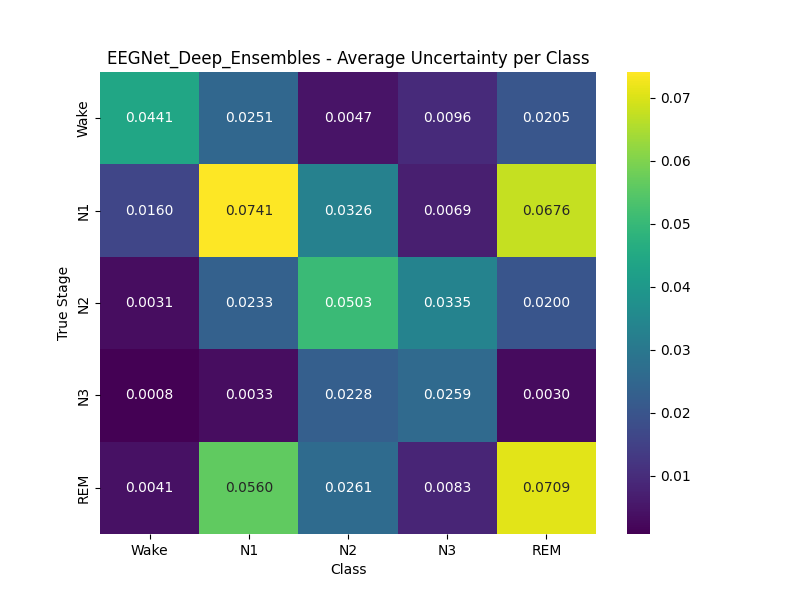

### uncertainty_heatmap_EEGNet_MC_Dropout.png

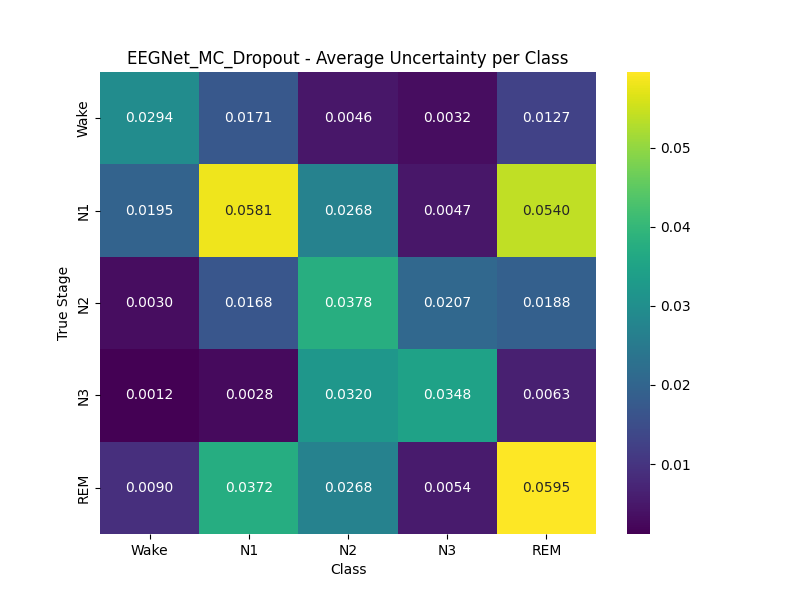

### uncertainty_heatmap_LSTM_Deep_Ensembles.png

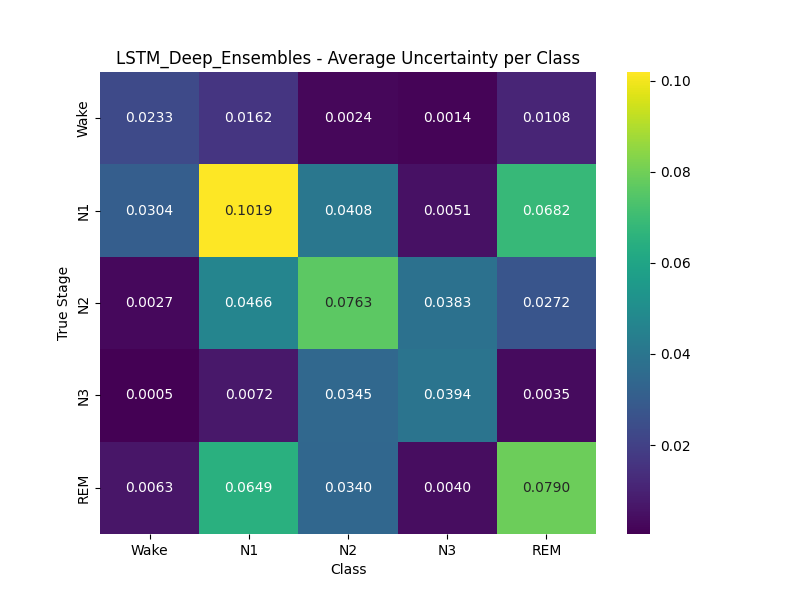

### uncertainty_heatmap_LSTM_MC_Dropout.png

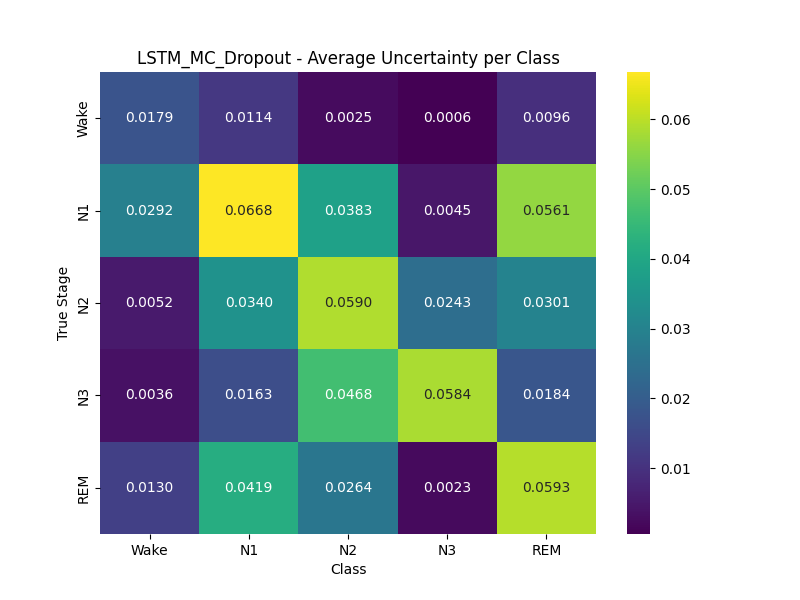

### uncertainty_violin_EEGNet_Deep_Ensembles.png

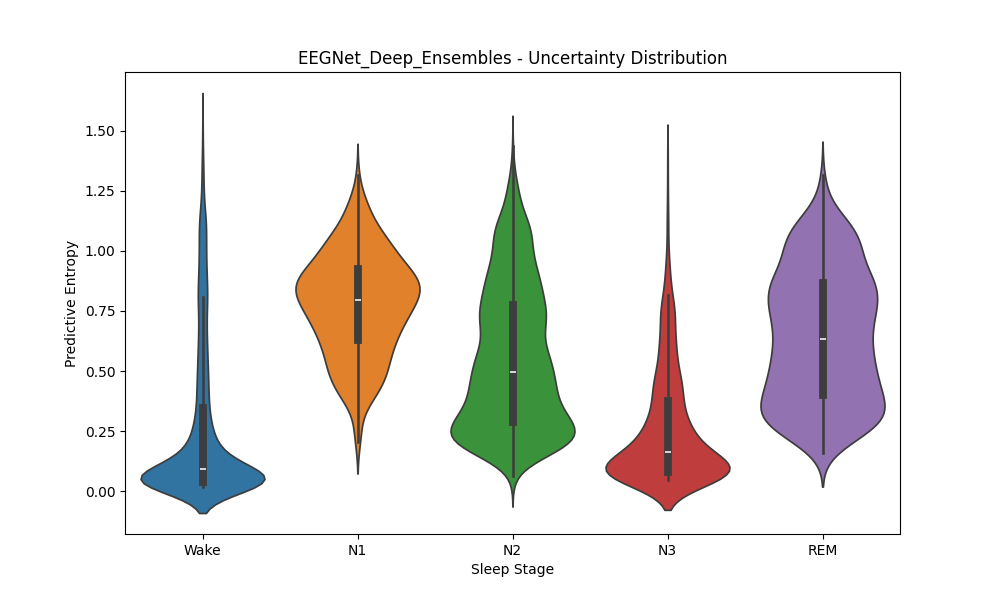

### uncertainty_violin_EEGNet_MC_Dropout.png

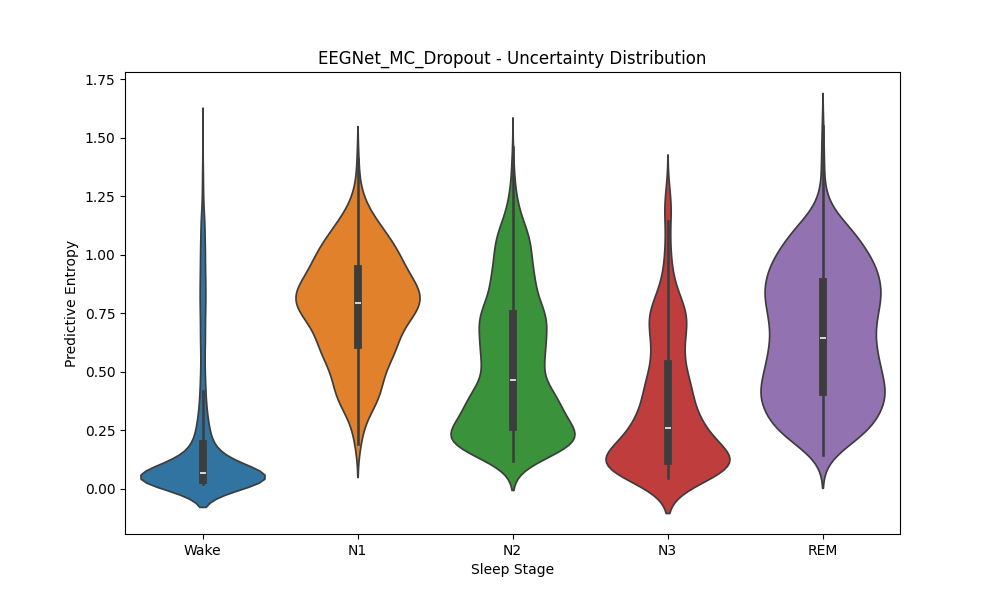

### uncertainty_violin_LSTM_Deep_Ensembles.png

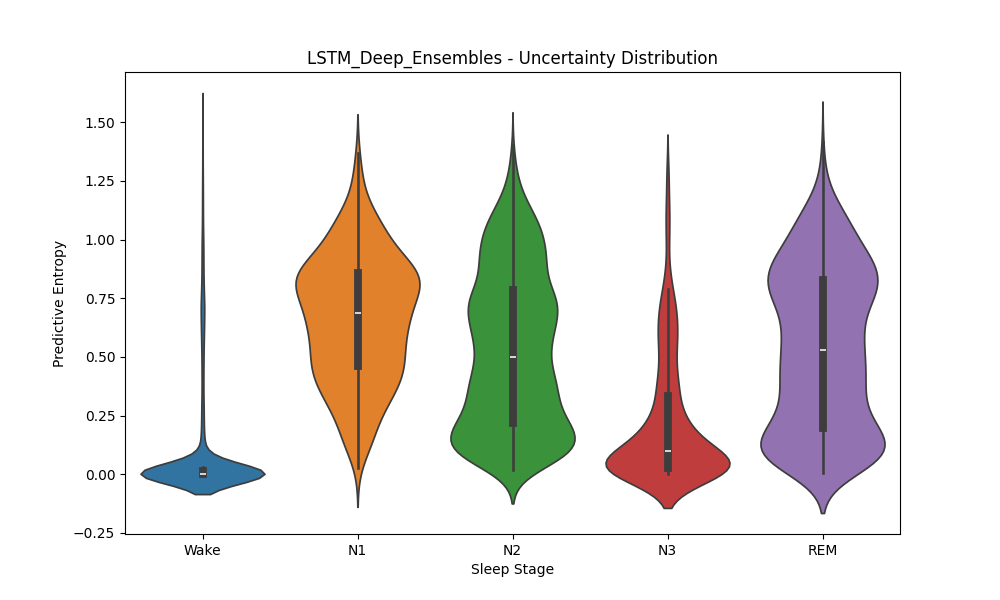

### uncertainty_violin_LSTM_MC_Dropout.png

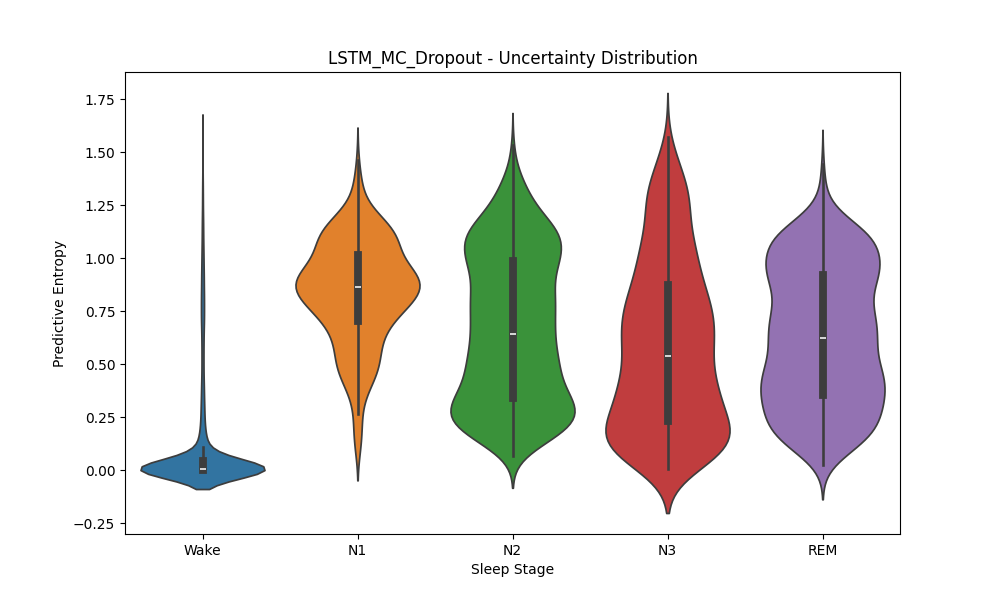
